# The *Tug1* Locus is Essential for Male Fertility

**DOI:** 10.1101/562066

**Authors:** Jordan P. Lewandowski, Gabrijela Dumbović, Audrey R. Watson, Taeyoung Hwang, Emily Jacobs-Palmer, Nydia Chang, Christian Much, Kyle Turner, Christopher Kirby, Jana Felicitas Schulz, Clara-Louisa Müller, Nimrod D. Rubinstein, Abigail F. Groff, Steve C. Liapis, Chiara Gerhardinger, Norbert Hubner, Sebastiaan van Heesch, Hopi E. Hoekstra, Martin Sauvageau, John L. Rinn

## Abstract

**Background:** Several long noncoding RNAs (lncRNAs) have been shown to function as central components of molecular machines that play fundamental roles in biology. While the number of annotated lncRNAs in mammalian genomes has greatly expanded, their functions remain largely uncharacterized. This is compounded by the fact that identifying lncRNA loci that have robust and reproducible phenotypes when mutated has been a challenge.

**Results:** We previously generated a cohort of 20 lncRNA loci knockout mice. Here, we extend our initial study and provide a more detailed analysis of the highly conserved lncRNA locus, Taurine Upregulated Gene 1 (*Tug1*). We report that *Tug1* knockout male mice are sterile with complete penetrance due to a low sperm count and abnormal sperm morphology. Having identified a lncRNA loci with a robust phenotype, we wanted to determine which, if any, potential elements contained in the *Tug1* genomic region (DNA, RNA, protein, or the act of transcription) have activity. Using engineered mouse models and cell-based assays, we provide evidence that the *Tug1* locus harbors three distinct regulatory activities – two noncoding and one coding: (i) a *cis* DNA repressor that regulates many neighboring genes, (ii) a lncRNA that can regulate genes by a *trans*-based function, and finally (iii) *Tug1* encodes an evolutionary conserved peptide that when overexpressed impacts mitochondrial membrane potential.

**Conclusions:** Our results reveal an essential role for the *Tug1* locus in male fertility and uncover three distinct regulatory activities in the *Tug1* locus, thus highlighting the complexity present at lncRNA loci.

## BACKGROUND

It has long been appreciated that noncoding RNAs play central roles in biology. Key cellular machines, such as telomerase and the ribosome, are comprised of both protein and noncoding RNAs and serve as classic examples of RNA-based functionalities (Feng et al., 1995; Sonenberg et al., 1975). LncRNAs have been shown to function in a variety of biological processes; however, different strategies to study lncRNA function have led to discrepancies in the observed phenotypes, thereby highlighting the challenges of finding robust and reproducible lncRNA phenotypes (Goudarzi et al., 2019). Moreover, another challenge presented when studying lncRNA loci, is that they can harbor several potential regulatory modalities including, DNA regulatory elements, misannotated protein-coding genes, and even the act of transcription. Therefore, it is important to determine what regulatory elements, if any, are active at lncRNA loci.

A number of studies have revealed that lncRNA loci can mediate their function through a variety of mechanisms (Kopp and Mendell, 2018). A few well-studied lncRNA examples include *Xist*, which is a key factor in the X inactivation pathway and acts locally (*cis*) (Lee and Jaenisch, 1997; Penny et al., 1996), *Malat1*, which modulates alternative splicing and acts distally (*trans*) (Tripathi et al., 2010), and other lncRNAs, such as *linc-Cox2*, which functions to activate and repress gene expression through local and distal mechanisms (Bester et al., 2018; Carpenter et al., 2013; Elling et al., 2018). While it is clear that a number of lncRNA loci have RNA-based roles, recent findings have shown that some lncRNA loci, such as *Lockd* (Paralkar et al., 2016), *lincRNA-p21* (Groff et al., 2016), and *Peril* (Groff et al., 2018), regulate gene expression in *cis* through DNA regulatory elements, independent of the noncoding transcript. Moreover, many lncRNAs possess small open reading frames (ORFs) (Housman and Ulitsky, 2016; Slavoff et al., 2013), and an increasing number encode small peptides that have biological roles (Anderson et al., 2015; Chng et al., 2013; Nelson et al., 2016; Stein et al., 2018). With this in mind, it is likely that more regulatory DNA, RNA and protein activities will be uncovered at lncRNA loci.

We previously reported the generation of 20 lncRNA loci knockout mouse strains, five of which displayed either viability, growth or brain phenotypes (Lai et al., 2015; Sauvageau et al., 2013). From the strains that did not initially display such phenotypes, we selected *Tug1* for further analysis because it is highly conserved between human and mouse and it has been reported to have a number of diverse cellular functions. *Tug1* was first identified to contain a lncRNA transcript that, upon RNAi-mediated knockdown, affects the development of photoreceptors in the mouse retina (Young et al., 2005). *Tug1* also has a human ortholog that has a number of unique molecular properties including being regulated by p53 (Guttman et al., 2009) and associating with polycomb repressive complex 2 (PRC2) (Khalil et al., 2009). In addition, *TUG1* RNA has also been proposed to play multiple cellular roles, such as acting as a tumor suppressor in human gliomas (Katsushima et al., 2016; Li et al., 2016), as a cytoplasmic miRNA sponge in prostate cancer cell lines (Du et al., 2016), and being involved in chromatin and gene regulation in the nucleus (He et al., 2018; Khalil et al., 2009; Long et al., 2016). Together, these studies highlight diverse cellular functions for the *Tug1* RNA.

Here, we characterize the *Tug1* locus using multiple genetic approaches and describe a physiological function in spermatogenesis and male fertility. We show that deletion of the *Tug1* locus in mice leads to male sterility due to reduced sperm production as well as a failure of spermatids to individualize during spermiation. Using several complementary genetic approaches (whole locus deletion with a *lacZ* reporter knock-in, an inducible *Tug1* transgene, and combinations thereof), we provide evidence of a DNA-based repressive element within the *Tug1* locus that regulates several genes in *cis*. Furthermore, we show that a gene-expression program dysregulated in *Tug1* knockout testes can be partially rescued by ectopic expression of *Tug1* RNA *in vivo*. Finally, we show that the *Tug1* locus contains an evolutionarily conserved ORF, which is translated into a peptide and regulates mitochondrial function upon overexpression. Collectively, our study implicates *Tug1* as an essential locus in male fertility and demonstrates that the *Tug1* locus contains at least three regulatory activities – two noncoding and one coding.

## RESULTS

### The *Tug1* lncRNA locus is widely expressed and highly conserved

The murine *Tug1* lncRNA locus is located on chromosome 11 and has three annotated transcripts (Figure 1A). *Tug1* shares a bidirectional promoter with its neighboring protein-coding gene *Morc2a*, whose transcription start site (TSS) is located approximately 680 base pairs upstream of the first *Tug1* TSS. The *Tug1* locus is enriched with hallmarks of active transcription, such as RNA polymerase II (Pol II) and histone H3 lysine 4-trimethylation (H3K4me3) at its promoter, H3K36me3 across its gene body, and abundant transcription as shown by RNA-seq (Figure 1A). However, the *Tug1* locus is simultaneously enriched with the repressive histone mark H3K9me3 in several mouse cell types (Figure 1A and Figure S1). This atypical combination of H3K9me3 and H3K36me3 histone marks at the *Tug1* locus is also conserved in human cells (Figure S1). Moreover, the binding of repressor proteins SIN3A and COREST has been detected at both the human and mouse promoters (Figure S1).

**Figure 1.**
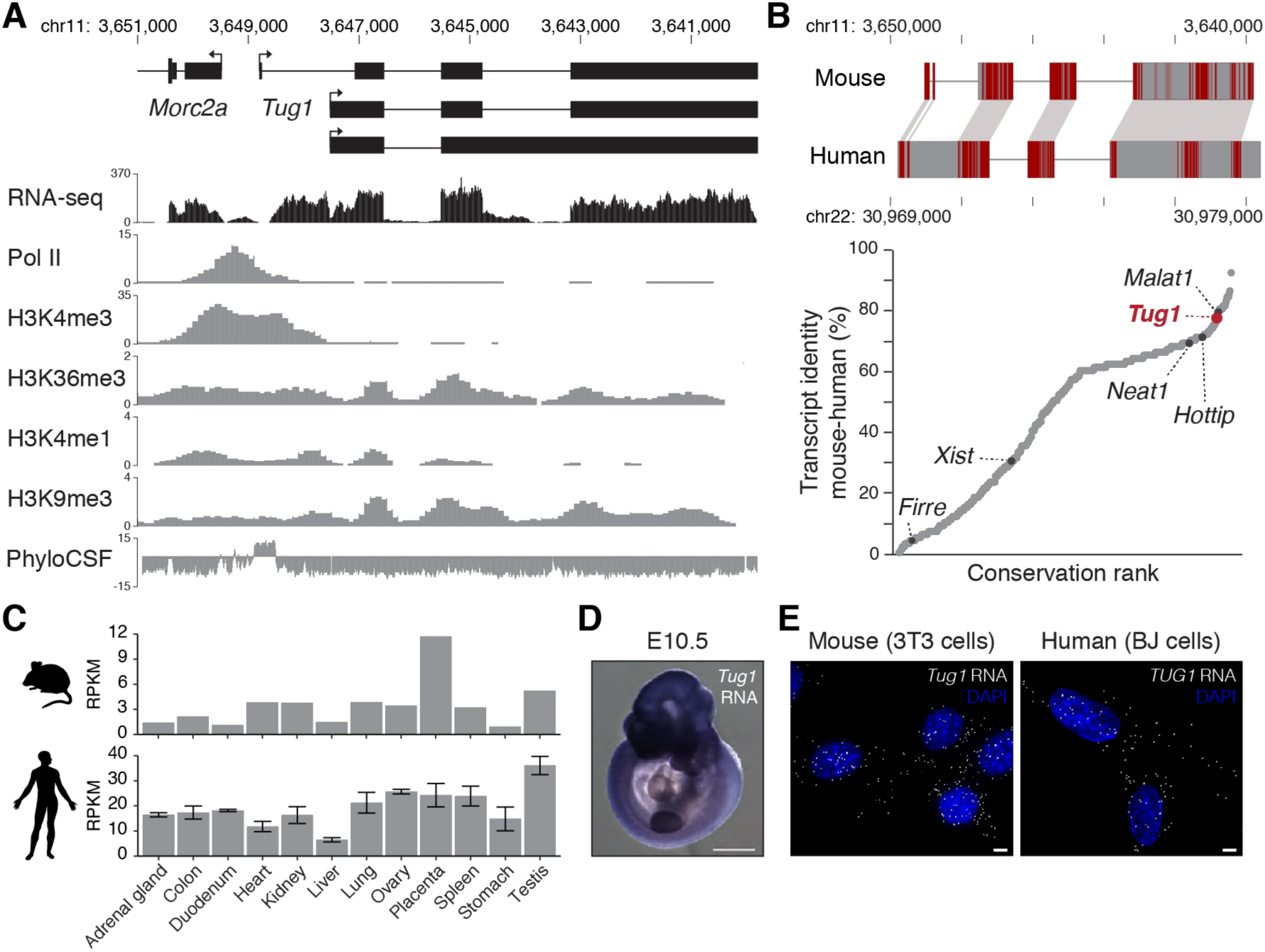
The *Tug1* lncRNA locus is highly conserved and ubiquitously expressed. **(A)** *Tug1* mouse genomic locus (shown inverted). UCSC Genome Browser tracks for RNA-sequencing (RNA-seq), RNA polymerase II (Pol II), histone 3 lysine 4-trimethylation (H3K4me3), H3K36me3, and H3K4me1 occupancy in testis and H3K9me3 occupancy in brain are depicted. PhyloCSF scores across the locus are shown. Chromosomal coordinates of the mouse *Tug1* gene are indicated (mm9). **(B)** Upper panel: Schematic of the nucleotide conservation alignment for mouse and human *Tug1*. Red lines indicate conserved nucleotides. Chromosomal coordinates of the *Tug1* gene for both species are indicated. Lower panel: Distribution of sequence identity for orthologous divergent and intergenic lncRNAs between mouse and human. X-axis shows increasing conservation rank. *Tug1* and other well characterized lncRNAs are highlighted. **(C)** RNA-seq expression levels of *Tug1* in a panel of mouse and human tissues. (**D**) RNA *in situ* hybridization of *Tug1* RNA in a mouse embryo at embryonic day 10.5 (E10.5). (**E**) Maximum intensity projections of *Tug1* single molecule RNA FISH (gray) on murine 3T3 and human BJ fibroblasts. Nucleus is stained with DAPI (blue). Scale bar is 5 μm.

*Tug1* is among the most conserved lncRNAs between human and mouse, with exonic nucleotide conservation levels reaching 77% (Figure 1B). This level of sequence conservation is similar to the highly abundant lncRNA *Malat1* (79%), and higher than other well characterized lncRNAs including *Hottip* (71%), *Neat1* (69%), *Xist* (30%) and *Firre* (4%) (Figure 1B) (Chen et al., 2016). Interestingly, further conservation analyses lead us to identify a highly conserved putative open reading frame (ORF) in the *Tug1 locus*, as indicated by phylogenic codon substitution frequencies (PhyloCSF) (Lin et al., 2011), a computational tool for identifying protein-coding and non-coding regions (Figure 1A).

Apart from its high level of sequence conservation, *Tug1* RNA also has unique expression properties. First, the *Tug1* lncRNA is expressed at moderate to high levels in several adult tissues in both mouse and human (Figure 1C) (Fagerberg et al., 2014; The Mouse ENCODE Consortium, 2014). Second, the *Tug1* lncRNA is abundantly detected in a number of embryonic tissues at different embryonic stages (E8.0 – E12.5) (Figure 1D and Figure S2). Finally, using single molecule RNA fluorescence *in situ* hybridization (smFISH) we observed that the *Tug1* lncRNA is detected in both the cytoplasm and the nucleus in human and mouse fibroblasts (Figure 1E), which is consistent with previous reports (Cabili et al., 2015; Khalil et al., 2009; van Heesch et al., 2014; Zhang et al., 2014).

### *Tug1*^-/-^ males are sterile due to impaired spermatogenesis

To investigate the *in vivo* role of *Tug1*, we utilized a previously generated full gene-ablation model (*Tug1*^*-/-*^), where after the promoter and first exon, the gene body of the *Tug1* locus was replaced with a *lacZ* reporter cassette, thereby keeping the act of transcription intact (Figure 2A) (Lai et al., 2015; Sauvageau et al., 2013). Notably, this deletion strategy also removed 86 out of 143 amino acids in the putative ORF (Figure S3). Loss of *Tug1* was confirmed by genotyping and by RNA-seq analysis in wild type and *Tug1*^*-/-*^ testes (Figure 2A). Thus, through this approach any potential phenotype due to the lncRNA, potential DNA elements or even the putative peptide would be included.

**Figure 2.**
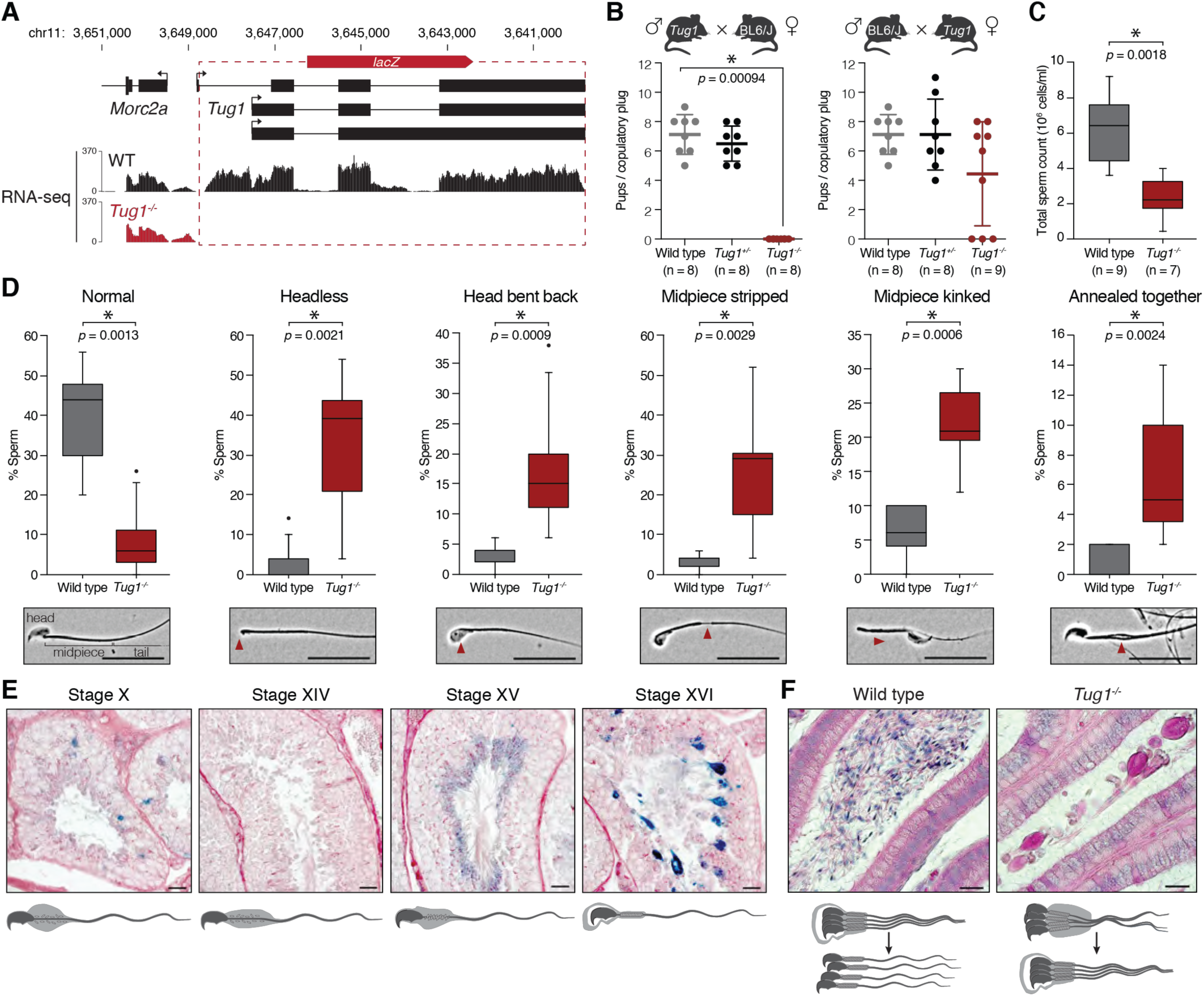
Deletion of the *Tug1* locus leads to sperm defects and male infertility. **(A)** Deletion strategy of the *Tug1* locus (shown inverted). The *Tug1* gene-body was replaced by a *lacZ* reporter cassette, leaving the promoter and first exon intact. The dashed lines indicate the deleted region in the *Tug1* knockout. RNA-sequencing (RNA-seq) tracks for wild type (WT) and *Tug1*^*-/-*^ testis are depicted. **(B)** Scatter dot plot (showing the mean and standard deviation) of the number of pups at birth per copulatory plug for matings between wild type, *Tug1*^*+/-*^ or *Tug1*^*-/-*^ males and wild type C57BL/6J females (left panel) and wild type C57BL/6J males and wild type, *Tug1*^*+/-*^ or *Tug1*^*-/-*^ females (right panel). Each dot represents a litter from a different mouse. Significant (*) *p*-value (Wilcoxon rank sum test with Bonferroni correction) and number of mice for each genotype tested are indicated. **(C)** Box plot of total sperm count for wild type and *Tug1*^*-/-*^ males. Significant (*) *p*-value (Wilcoxon rank sum test) is indicated. **(D)** Box plots of the percentage of normal sperm and sperm with the five most common morphological abnormalities for wild type (n = 9) and *Tug1*^*-/-*^ (n = 8) males. Representative images of normal and morphologically aberrant sperm are located below each corresponding plot. Red arrows indicate the location of the defect. Scale bars are 20 μm. Significant (*) *p*-values (Wilcoxon rank sum test) are indicated. **(E)** Representative spermatocyte diagrams and micrographs of *Tug1*^*+/-*^ seminiferous tubule sections stained with periodic acid-Schiff’s reagent and X-gal showing expression of the *lacZ* reporter under the control of the endogenous *Tug1* promoter at the indicated stages of spermatogenesis. Scale bars are 20 μm. **(F)** Representative spermatid diagrams and micrographs of wild type and *Tug1*^*-/-*^epididymis tubule sections stained with hematoxylin and eosin. Scale bars are 20 μm.

*Tug1*^-/-^ mice are viable and do not display any obvious physiological abnormalities up to one year of age, with the exception of a slight reduction in weight in male mice relative to wild type littermates (Figure S4A). As previously reported, the progeny of *Tug1*^*+/-*^ intercrosses follow normal Mendelian ratios (Sauvageau et al., 2013). However, we noticed a complete absence of offspring from intercrosses between *Tug1*^*-/-*^ mice (n = 4 breeding pairs). Therefore, we sought to investigate the fertility of *Tug1*^-/-^ mutants in more detail. We separately mated *Tug1*^*-/-*^, *Tug1*^*+/-*^ and wild type males or females to *C57BL/6J* mice. We did not observe a difference in the mounting behavior between wild type and *Tug1*^-/-^ mice, as assessed by the presence of a vaginal plug. Strikingly, matings between *Tug1*^*-/-*^ males (n = 8) and *C57BL/6J* females did not produce any offspring, whereas matings involving either *Tug1*^*+/-*^ males (n = 8) or wild type males (n = 8) with *C57BL/6J* females resulted in similar numbers of offspring (Figure 2B). Moreover, six out of nine *Tug1*^*-*^*/-* females that mated with *C57BL/6J* males gave birth to pups (Figure 2B), indicating that only *Tug1*^*-/-*^ males appear sterile. Thus, the *Tug1* locus is likely required for male fertility.

To further understand the underlying fertility defect in *Tug1*^-/-^ males, we first examined the reproductive morphology of wild type and *Tug1*^-/-^ male mice. Testicular descent appeared normal and we did not observe any other gross morphological abnormalities in their reproductive system upon dissection (Figure S4B). We measured testes mass relative to total body weight and did not observe a significant decrease (*p* = 0.0751) in *Tug1*^-/-^ (mean = 0.25 ± 0.020 %, n = 8) compared to wild type (mean = 0.30 ± 0.016 %, n = 9) (Figure S4C). Next, we quantified sperm production and found a significant reduction in sperm number from *Tug1*^-/-^ males (mean = 2.35 x 10^6^ ± 0.473 x 10^6^ cells/mL, n = 7), which produced on average only 40% as many sperm as wild type mice (6.13 x 10^6^ ± 0.636 x 10^6^ cells/mL, n = 9, *p* = 0.0018) (Figure 2C). Notably, although *Tug1*^-/-^ males produce fewer sperm, none were found to completely lack sperm (azoospermic).

Based on these results, we investigated whether perturbations in sperm morphology could explain the complete infertility in *Tug1*^-/-^ males. We examined the morphological features of sperm and quantified the frequency of 15 different abnormalities (Table S1). Overall, the proportion of morphologically normal sperm was significantly lower in *Tug1*^*-/-*^ mice (mean = 8.3 ± 3.0 %, n = 8, *p* = 0.0013) compared to wild type males (mean = 38.9 ± 4.3 %, n = 9) (Figure 2D). We observed significant morphological defects in *Tug1*^-/-^ sperm including: sperm with no head, misshapen head, head bent back, stripped midpiece, kinked midpiece, curled midpiece, midpiece debris, broken tail, and the presence of multiple sperm attached along the midpiece (Figure 2D, Figure S4D, and Table S1). Together, these results indicate that the sterility of *Tug1*^-/-^ males arises from a combination of low sperm count (oligozoospermia) and abnormal sperm morphology (teratozoospermia).

To further investigate how the deletion of the *Tug1* locus leads to abnormal sperm morphology, we examined the timing of *Tug1* expression at different stages of spermatogenesis. To this end, we took advantage of the knock-in *lacZ* reporter driven by the endogenous *Tug1* promoter and assessed expression by *lacZ* staining of histological sections of *Tug1*^*+/-*^ testis and epididymis. From stages IX to XI of spermatogenesis in the testis, *lacZ* staining was restricted to excess cytoplasm, known as residual bodies, which are phagocytosed toward the basement membrane by Sertoli cells (Figure 2E) (Firlit and Davis, 1965). No expression was detected in the later stages XII to XIV (Figure 2E). However, we observed *lacZ* staining in stage XV elongated spermatids and the *lacZ* staining became stronger at stage XVI, just before spermiation (Figure 2E). The observed *lacZ* pattern indicates that *Tug1* expression is temporally controlled during spermatogenesis.

In *Tug1*^-/-^ testes, mature spermatids appeared to remain attached by their collective cytoplasm. This was even more striking in the epididymis, where multiple sperm aggregates were observed in *Tug1*^-/-^ mice, while individual sperm appeared to migrate freely throughout the lumen in wild type mice (Figure 2F). These aggregates were present in all regions of the epididymis (caput, corpus and cauda). Consistent with the reduced sperm count, fewer individual sperm were observed in *Tug1*^-/-^ epididymis tissue compared to wild type. Together, our analyses of the *Tug1*^-/-^ model provide evidence that the locus is required for male fertility.

### *Tug1* DNA encodes a *cis* repressor regulatory element

Since we observed a robust phenotype in our *Tug1*^-/-^ model, we next sought to investigate what, if any, molecular activities (DNA, lncRNA, and protein) are present at the *Tug1* locus. We first focused on determining if the DNA at the *Tug1* locus harbored any regulatory activity, because many lncRNA loci have been reported to contain DNA regulatory elements that can regulate the expression of neighboring genes (*cis*-acting) (Groff et al., 2016; Groff et al., 2018; Paralkar et al., 2016). Our *Tug1*^-/-^ model enables us to test for potential *cis* regulatory activity within the *Tug1* locus because our gene-ablation design removes potential *cis*-acting elements, yet keeps the act of transcription intact (Figure 2A) (Lai et al., 2015; Sauvageau et al., 2013). To determine if there is a local regulatory effect on gene expression, we performed RNA-seq on testes from wild type and *Tug1*^-/-^ mice and plotted significant changes in gene expression within a 2-Mb region centered on the *Tug1* locus (FDR < 0.05, FC > 1.5). Of the 71 genes within this window, we observed six differentially regulated genes: *Rnf185, Pla2g3, Selm, Smtn, Gm11946* and *8430429K09Rik*. Notably, all of these genes were significantly upregulated in *Tug1*^-/-^ compared to wild type and located downstream of the *Tug1* TSS (Figure 3A). Because these six genes are all upregulated in *Tug1*^-/-^ testes, this local effect on neighboring gene expression provides evidence of a *cis* repressor function in the *Tug1* locus.

**Figure 3.**
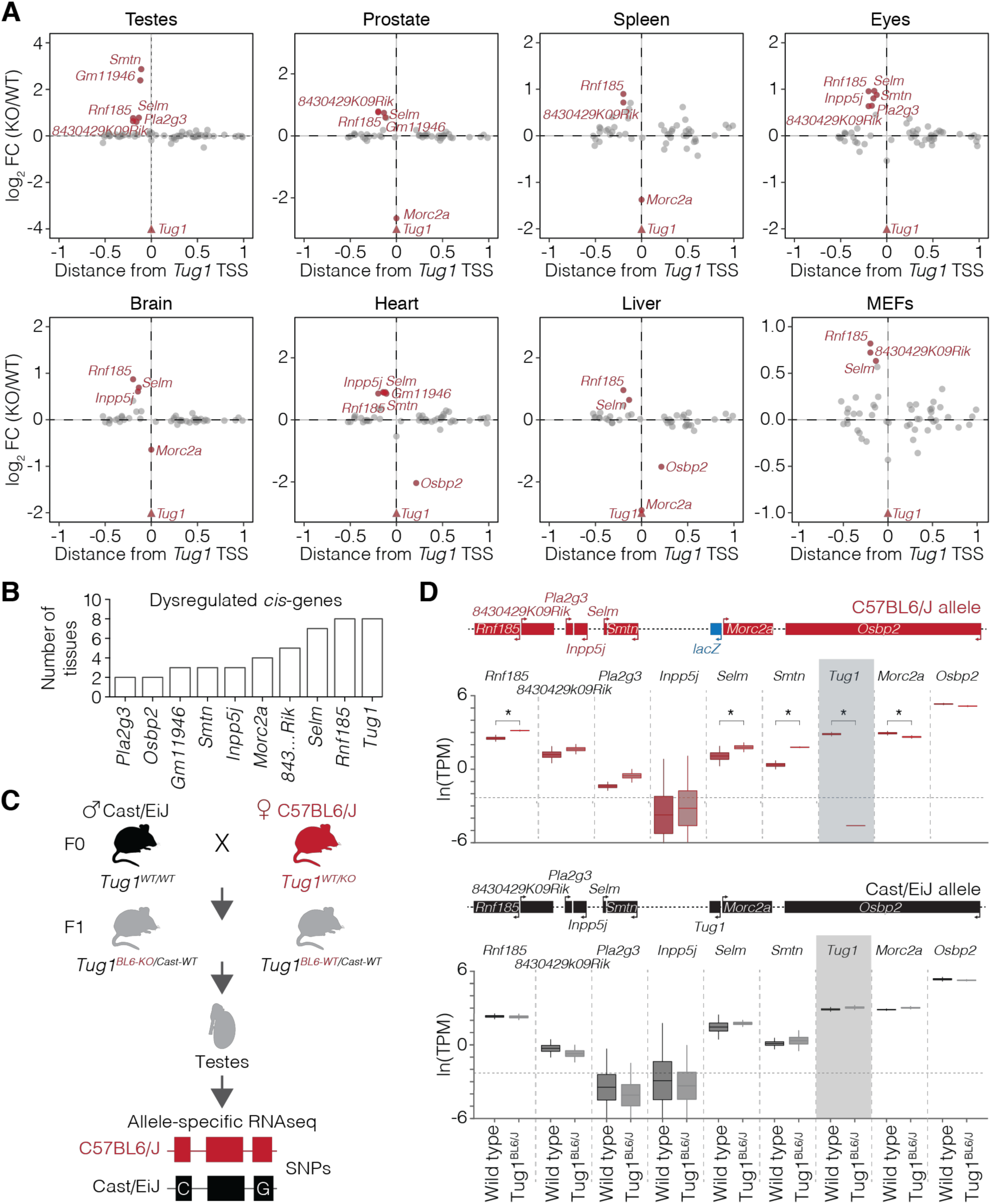
The *Tug1* locus harbors a *cis*-repressive DNA regulatory element. **(A)** Differential expression of genes in the local region (±1 Mb) of *Tug1* for each indicated mouse tissue, depicted as fold change (FC) between *Tug1*^*-/-*^ (KO) and wild type (WT). Significantly differentially expressed genes are marked and labeled in red. **(B)** Plot of the number of tissues that genes downstream of *Tug1* TSS are found significantly dysregulated. **(C)** Strategy for allele-specific RNA-sequencing (RNA-seq). *Tug1*^*+/-*^ C57BL/6J females were crossed with wild type Cast/EiJ males and testes from the F1 hybrid progeny were harvested for RNA-seq. Single nucleotide polymorphisms (SNPs) allowed for the differentiation between the C57BL/6J and the Cast/EiJ allele. **(D)** Allele-specific expression of local genes surrounding *Tug1* in testes from F1 hybrid C57BL/6J∷Cast/EiJ wild type (*Tug1*^*BL6-WT/Cast-WT*^) and heterozygous *Tug1* knockout (*Tug1*^*BL6-KO/Cast-WT*^) mice containing a deletion on the C57BL/6J allele. Upper panel, expression levels of neighboring genes from the C57BL/6J allele; lower panel, expression levels of neighboring genes from the Cast/EiJ allele. Boxes are centered at the mean, extend one standard deviation, and the bottom and top notches are the minimum and maximum samples. The genomic locus encompassing the local genes around *Tug1* is depicted. Asterisks indicate significant Bayesian posterior probability (PP>0.95) differential expression between hybrid wild type and *Tug1*^*BL6-KO*^testes. Horizontal dotted line indicates expression levels below 0.1 TPM.

To further investigate whether the *cis*-effect of the *Tug1* locus was more widespread, we performed RNA-seq on six additional tissues (prostate, spleen, eyes, heart, liver and mouse embryonic fibroblasts (MEFs)) as well as re-analyzed an existing brain dataset (Goff et al., 2015) from wild type and *Tug1*^-/-^ mice (Table S2). We examined whether genes within a 2-Mb window centered on the *Tug1* locus were similarly dysregulated in the different tissues. Consistent with the testes, of the 71 genes within this window, nine genes were dysregulated in one or more tissues (seven upregulated and two downregulated) (Figure 3A). Notably, of the seven upregulated genes, the E3 ubiquitin ligase *Rnf185* was consistently upregulated in 8 of 8 *Tug1*^-/-^ tissues, followed by the selenoprotein gene, *Selm* (7 of 8 samples), and *8430429K09Rik* (6 of 8 samples) (Figure 3B). This dysregulation is consistent with a previous study from our group in which we observed a misregulation of genes located near the *Tug1* locus in the brain of our *Tug1*^-/-^ model (Goff et al., 2015). We also observed that *Morc2a*, the protein-coding gene that shares a promoter with *Tug1*, was significantly downregulated in 4 of the 8 samples. Collectively, these data suggest that the *Tug1*-mediated repressive *cis*-effect functions in a broad range of tissues.

Since the neighboring genes are upregulated upon deletion of the *Tug1* locus, we reasoned that the repressive activity could be mediated either directly by the *Tug1* transcript or by regulatory DNA elements within the locus. To determine if the repressive effect of *Tug1* on neighboring genes occurs on the same allele (*cis-*acting), we performed allele-specific RNA-seq using a hybrid mouse strain. To generate this strain, we crossed *Tug1*^*+/-*^ C57BL/6J females with *Mus castaneus* (Cast/EiJ) males (Figure 3C). The resulting polymorphisms in the F1 hybrid progeny (∼1/150 bp between C57BL/6J and Cast/EiJ) allow quantification of gene expression from each strain-specific allele (Keane et al., 2011). We thus harvested testes from F1 hybrid males harboring a maternal C57BL/6J allele deletion and performed allele-specific expression analysis (Figure 3B and Table S3). As a control for haplotype specific effects, we also analyzed allele-specific expression differences in wild type F1 hybrid C57BL/6J∷Cast/EiJ male littermates. We then quantified the expression from each allele and found that *Rnf185, Selm*, and *Smtn* were significantly upregulated and *Morc2a* slightly downregulated only on the C57BL/6J allele containing the *Tug1* deletion (Figure 3D). Importantly, no change in expression was detected from any gene within 1 Mb of *Tug1* on the Cast/EiJ allele, which contains an intact *Tug1* locus (Figure 3D). Moreover, it is notable that *Tug1* RNA from the intact allele does not impact the dysregulated genes found on the *Tug1* knockout allele, thereby suggesting a DNA-based repressor role at the *Tug1* locus. From the multiple mouse models, we conclude that the *Tug1* DNA, rather than the lncRNA or the act of transcription, exerts a repressive effect in *cis* on several genes up to 200 kb downstream of the *Tug1* transcription site.

### *Tug1* lncRNA regulates gene expression in *trans*

Previous studies have suggested a *trans* role for the *Tug1* lncRNA on chromatin regulation and gene expression (Han et al., 2013; Li et al., 2016; Long et al., 2016; Xu et al., 2014; Young et al., 2005; Zhang et al., 2014). Thus, we set out to determine if the lncRNA from the *Tug1* locus displays any trans regulatory activity on gene expression *in vivo*. We analyzed the RNA-seq data for *Tug1*^-/-^ tissues (testis, prostate, spleen, eyes, liver, heart, brain and MEFs), and identified significant changes in gene expression relative to wild type. Deletion of the *Tug1* locus was accompanied by 2139 significantly dysregulated genes across all tissues examined. We observed that global changes in gene expression clustered by tissue-type, indicating tissue-specific gene dysregulation (Figure 4A, Table S2, and Table S4). We found that while most dysregulated genes (∼89%) were perturbed in only a single tissue (Figure 4B), several genes were commonly dysregulated across multiple tissues (Figure 4B, Table S2, Table S4). We then performed gene set enrichment analysis (GSEA) using the differentially expressed genes for each tissue and observed an enrichment of several pathways that were shared across the individual tissues. For example, oxidative phosphorylation, Myc targets, and epithelial to mesenchymal transition were found enriched in 7 of the 8 *Tug1*^-/-^ tissues (Figure 4C).

**Figure 4.**
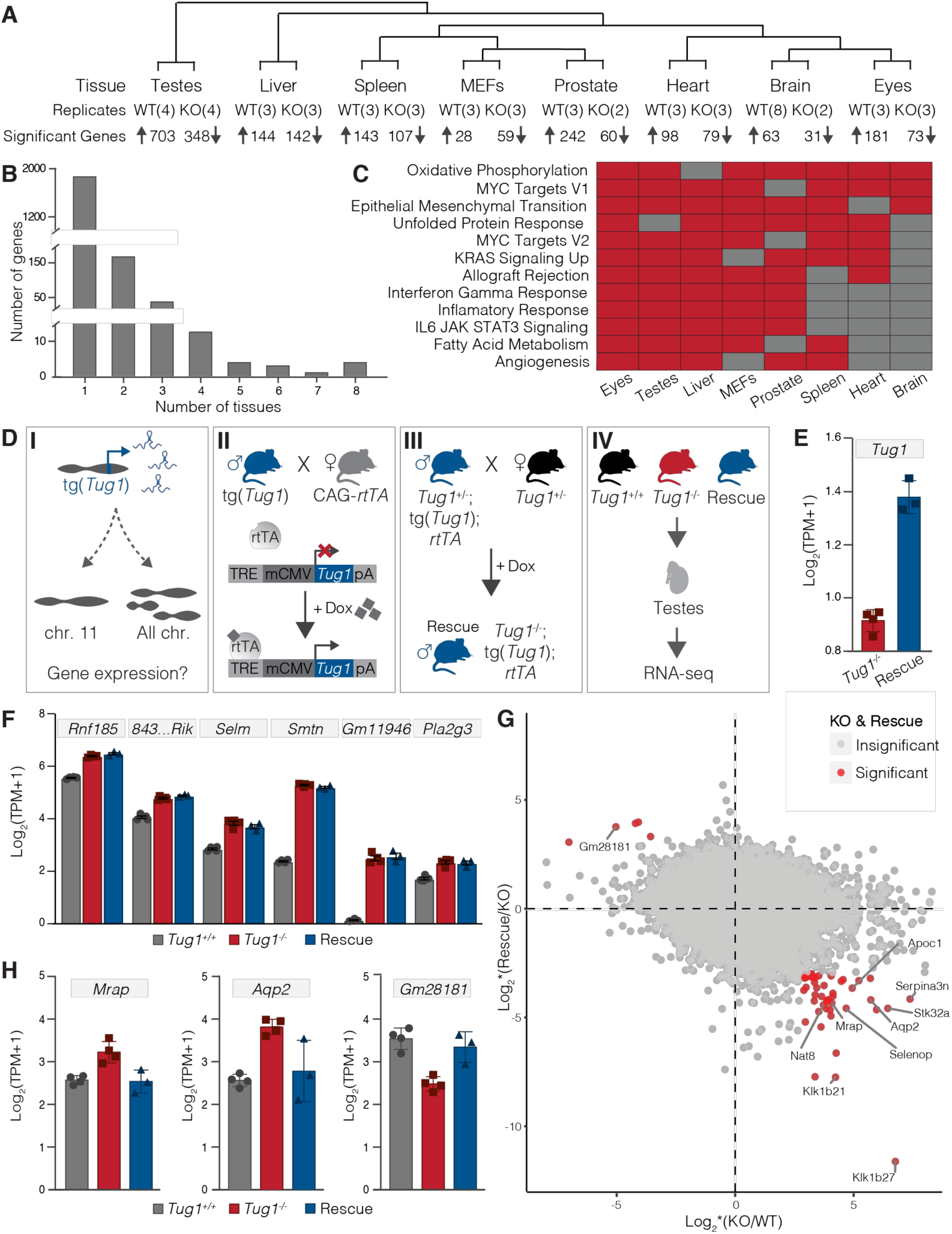
*Tug1* lncRNA regulates gene expression in *trans.* **(A)** Adult tissue types and mouse embryonic fibroblasts (MEFs) used for RNA sequencing of wild type (WT) and *Tug1*^*-/-*^ (KO) mice. For each tissue, the number of biological replicates per genotype and the number of upregulated and downregulated genes (FDR<0.05) is shown from KO to WT comparisons. The number of perturbed genes (y-axis) in KO animals according to the number of tissues in which the gene was found to be dysregulated (x-axis). **(C)** Gene Set Enrichment Analysis (GSEA) of differentially expressed genes found in wild type vs. *Tug1*^-/-^ murine tissues and MEFs. Red shading indicates tissue in which gene set is perturbed, grey shading indicates tissue in which gene set is not different between WT and KO. **(D)** Schematic showing the experimental design to identify genes reciprocally regulated by *Tug1* RNA. **(DI)** Testing the impact of the *Tug1* transgene expression on gene expression *in vivo* **(DII)** Schematic of the *Tug1* transgene (tg(*Tug1*)) and systemic induction by mating to CAG-rtTA3 mice in the presence of doxycycline (dox). **(DIII)** Schematic of matings to generate *Tug1*^rescue^ mice (*Tug1*^-/-^; tg(*Tug1*); *rtTA*), enabling dox-inducible *Tug1* expression in a *Tug1*-knockout background. **(DIV)** Collection of testes from WT (*Tug1*^+/+^) (n = 4), KO (*Tug1*^*-/-*^) (n = 4), and *Tug1*^rescue^ (n = 3) mice for RNA sequencing. **(E)** *Tug1* gene expression level (log_2_TPM+1) in testes of KO (red) and doxycycline-induced *Tug1*^rescue^ (blue) mice. Error bars represent standard error of the mean. **(F)** Expression levels (log_2_TPM+1) for *Tug1* neighboring genes in WT (grey), KO (red), and *Tug1*^rescue^ (blue) mice. **(G)** Genome-wide profile of reciprocally regulated genes from *Tug1* RNA induction. The fold change score of KO-WT is plotted on the x-axis and the fold change score of Rescue-KO is plotted on the y-axis. The fold change score (*) is the fold change divided by standard deviation. Genes significantly differentially regulated in both comparisons of KO-WT and Rescue-KO are labeled in red, otherwise labeled in grey. Examples of reciprocally regulated genes are labeled with the gene name. **(H)** Examples of differentially expressed genes in testes showing significant reciprocal regulation in WT (grey), KO (red), and *Tug1*^rescue^ (blue) mice.

To investigate the role of *Tug1* RNA, we sought to address whether ectopic expression of *Tug1* RNA could restore the genes dysregulated in *Tug1*^-/-^ testes. Given that *Tug1* harbors a putative peptide encoded in the 5’ region (discussed below), we focused on a *Tug1* isoform that lacks the 5’ region, thus ensuring we would address the role of *Tug1* RNA alone. To this end, we generated a doxycycline (dox)-inducible *Tug1* transgenic mouse by cloning a *Tug1* isoform downstream of a tet-responsive element (henceforth called tg(*Tug1*)) (Figure 4D and methods section). Next, we generated compound transgenic mice that contained the *Tug1* transgene in the *Tug1*^-/-^ background that also constitutively overexpressed the reverse tetracycline transcriptional activator gene (CAG-rtTA3) (combined alleles henceforth called *Tug1*^rescue^). This approach enabled systemic induction of *Tug1* RNA in the presence of dox, allowing to distinguish DNA- and RNA-based effects, and to test if *Tug1* RNA expression alone would be sufficient to rescue gene expression and male fertility phenotypes arising in *Tug1*^-/-^ mice.

Because *Tug1*^rescue^ mice lacked endogenous *Tug1*, we were able to assess the level of *Tug1* RNA from the transgene. We performed RNA-seq on testes from *Tug1*^rescue^ mice (*n* = 3) and found that *Tug1* RNA from the transgene was expressed at significantly lower levels than wild type in the testes (Figure 4E and Figure S5A). Moreover, we sorted peripheral blood cell types (CD4, CD8, and NK) from *Tug1*^rescue^ mice and also found lower levels of *Tug1* RNA induction relative to wild type (Figure S5A,B). Even though the transgene expression was low, we reasoned that this would still be a valuable *in vivo* model to test RNA-mediated effects on gene regulation. Thus, we tested whether genes found dysregulated in the testes from *Tug1*^-/-^ mice could be rescued by ectopic expression of the *Tug1* RNA in our *Tug1*^rescue^ model. Notably, 52 of the 1051 genes that were dysregulated in *Tug1*^-/-^ testes were found significantly reciprocally regulated in *Tug1*^rescue^ testes (Figure 4G, Table 1, and Table S4). For example, a mitochondrial related gene, *Mrarp*, and an aquaporin gene, *Aqp2*, are significantly upregulated in *Tug1*^-/-^ testes, but their expression was reduced to wild type levels in *Tug1*^rescue^ testes (Figure 4H). Conversely, the predicted lncRNA gene *Gm28181* that is significantly reduced in *Tug1*^-/-^ testes, is significantly upregulated to wild type levels in *Tug1*^rescue^ testes (Figure 4H). While we observed a *trans*-effect for *Tug1* RNA, we did not observe any changes in expression for the genes neighboring the *Tug1* locus (Figure 4F and Table S4). Taken together, these data demonstrate that *Tug1* lncRNA regulates a subset of genes by an RNA-based *trans* mechanism, evident even at low levels of *Tug1* RNA.

**Table 1.** Genes reciprocally regulated by *Tug1* lncRNA in the testes. List of genes with significant changes in expression between wild type (WT), *Tug1*^-/-^ (KO), and *Tug1*^rescue^ testes. Chromosomal location of the genes, mean TPM for each condition, and the main biological processes associated with each gene are listed. Significance of the fold change between wild type verses *Tug1*^-/-^ (WT-KO) and *Tug1*^-/-^ verses *Tug1*^rescue^ (KO-Rescue) is indicated by asterisks (*, p<0.05; **, p<0.01, ***, p<0.001).

| Gene name | Location | Mean TPM |  |  | Significance |  | Biological process |
| --- | --- | --- | --- | --- | --- | --- | --- |
|  |  | WT | KO | Rescue | WT-KO | KO-Rescue |  |
| *Pard3b* | chr1 | 2.66 | 2.13 | 2.74 | *** | *** | microtubule cytoskeleton organization |
| *Nav1* | chr1 | 1.40 | 2.33 | 1.45 | ** | ** | microtubule bundle formation |
| *Gm28181* | chr1 | 10.81 | 4.63 | 9.39 | *** | ** | unknown |
| *Col5a1* | chr2 | 7.16 | 8.97 | 7.85 | * | * | cell adhesion |
| *Hoxd10* | chr2 | 2.17 | 4.43 | 2.23 | * | * | regulation of transcription |
| *Sulf2* | chr2 | 12.20 | 16.74 | 13.21 | *** | ** | metabolic process |
| *Amy1* | chr3 | 16.43 | 23.06 | 14.58 | * | *** | metabolic process |
| *Dhcr24* | chr4 | 30.92 | 43.38 | 34.72 | *** | * | lipid metabolic process |
| *Tmem176a* | chr6 | 17.89 | 25.23 | 16.32 | ** | *** | lipid metabolic process |
| *Nat8f5* | chr6 | 2.30 | 4.78 | 2.13 | ** | ** | system development |
| *Nat8* | chr6 | 9.57 | 16.53 | 8.23 | ** | *** | glutathione metabolic process |
| *Apoc1* | chr7 | 35.49 | 56.71 | 37.27 | *** | ** | lipid metabolic process |
| *Vmn1r181* | chr7 | 2.65 | 4.26 | 1.88 | ** | *** | unknown |
| *Klk1b27* | chr7 | 12.21 | 31.01 | 2.65 | *** | *** | proteolysis |
| *Klk1b21* | chr7 | 23.81 | 56.40 | 5.45 | *** | *** | proteolysis |
| *Klk1b24* | chr7 | 19.20 | 34.33 | 5.95 | * | *** | proteolysis |
| *Cd209f* | chr8 | 1.57 | 2.64 | 1.23 | * | *** | cell adhesion |
| *Gpt2* | chr8 | 25.46 | 33.70 | 24.06 | ** | *** | biosynthetic process |
| *Tug1* | chr11 | 40.08 | 0.89 | 1.60 | *** | *** | - |
| *Spns2* | chr11 | 6.30 | 7.92 | 6.47 | * | * | sphingolipid metabolic process |
| *Serpina3n* | chr12 | 4.10 | 13.85 | 7.18 | *** | *** | inflammatory response |
| *Ankrd9* | chr12 | 43.02 | 50.58 | 44.74 | * | * | post-translational protein modification |
| *Sv2c* | chr13 | 5.59 | 7.53 | 5.15 | *** | *** | transmembrane transport |
| *3110070M22Rik* | chr13 | 110.87 | 93.69 | 120.25 | ** | * | unknown |
| *Tmem267* | chr13 | 87.89 | 77.92 | 92.25 | *** | *** | unknown |
| *Il17rb* | chr14 | 4.20 | 6.08 | 3.65 | *** | *** | regulation of cell growth |
| *Stab1* | chr14 | 3.99 | 5.50 | 4.04 | ** | * | cell adhesion |
| *Selenop* | chr15 | 88.40 | 130.97 | 85.90 | *** | *** | selenium compound metabolic process |
| *C7* | chr15 | 94.22 | 128.27 | 91.48 | ** | *** | immune response |
| *Aqp2* | chr15 | 4.93 | 13.17 | 6.48 | *** | *** | water transport |
| *AU021092* | chr16 | 30.38 | 44.08 | 33.37 | *** | * | unknown |
| *Nrros* | chr16 | 3.00 | 3.93 | 2.99 | * | * | superoxide metabolic process |
| *Mrap* | chr16 | 4.83 | 8.21 | 4.75 | *** | *** | protein localization to plasma membrane |
| *Cyp21a1* | chr17 | 1.99 | 3.12 | 1.22 | * | *** | steroid metabolic process |
| *Ston1* | chr17 | 28.51 | 34.85 | 29.11 | *** | *** | regulation of endocytosis |
| *Stk32a* | chr18 | 1.22 | 2.78 | 1.50 | *** | *** | protein phosphorylation |
| *mt-Rnr1* | chrM | 211.26 | 253.11 | 219.24 | * | * | ribosome biogenesis |

We also tested if *Tug1*^rescue^ male mice had normal fertility. We did not obtain any progeny from matings between *Tug1*^rescue^ male mice (*n* = 3) with C57BL6/J female mice (*n* = 12) (Figure S5C). Moreover, we found that *Tug1*^rescue^ males had a low sperm count (mean = 3.20 x 10^5^ ± 8.0 x 10^3^ cells/mL) which was similar to the lower sperm count observed in *Tug1*^-/-^ males (mean = 4.69 x 10^5^ ± 1.6 x 10^4^ cells/mL) compared to wild type (mean = 9.32 x 10^5^ ± 3.9 x 10^3^ cells/mL). In addition, histological sections of *Tug1*^rescue^ testes and epididymis showed fewer sperm, thereby confirming the low sperm count (Figure S5E). In further analysis, we observed that *Tug1*^rescue^ mice had a low proportion of normal shaped sperm which was also observed in *Tug1*^-/-^ mice (Figure S5F). While this finding may indicate that the sterility phenotype is not due to the lncRNA, the lack of a fertility rescue may also be due to the insufficient levels of *Tug1* expression from the transgene in the testes.

### The *Tug1* locus encodes an evolutionary conserved peptide in human and mouse

It has become increasingly clear that some lncRNA annotations also encode small peptides (Makarewich and Olson, 2017). Since a PhyloCSF analysis revealed the presence of putative ORFs in the *Tug1* locus (Figure 1A and Figure S6A), we further tested whether the *Tug1* locus could encode a peptide using biochemical and cell-based assays. First, we systematically screened for ORFs that displayed strong conservation across species, allowing for both canonical (AUG) and non-canonical (CUG and UUG) translation start codons. We identified multiple short ORFs in human and mouse *TUG1*/*Tug1* (11 and 15, respectively) (Figure 5A). Two ORFs (designated as ORF1 and ORF2) at the 5’ region of *TUG1*/*Tug1* drew our attention due to their conserved translational start and stop sites, as well as their high level of nucleotide conservation between human and mouse (Figure 5A). ORF1 (154 amino acids in human) and ORF2 (153 amino acids in human) both start with a non-canonical start codon (CUG). On the amino acid level, ORF1 and ORF2 share 92% and 70% cross-species identity, respectively. Moreover, ORF1 has a high PhyloCSF score (350) and shows conservation spanning its entire sequence, whereas ORF2 does not show patterns of preserving synonymous mutations, indicating that ORF1 is more likely to be translated (Figure 5B and Figure S6A).

**Figure 5.**
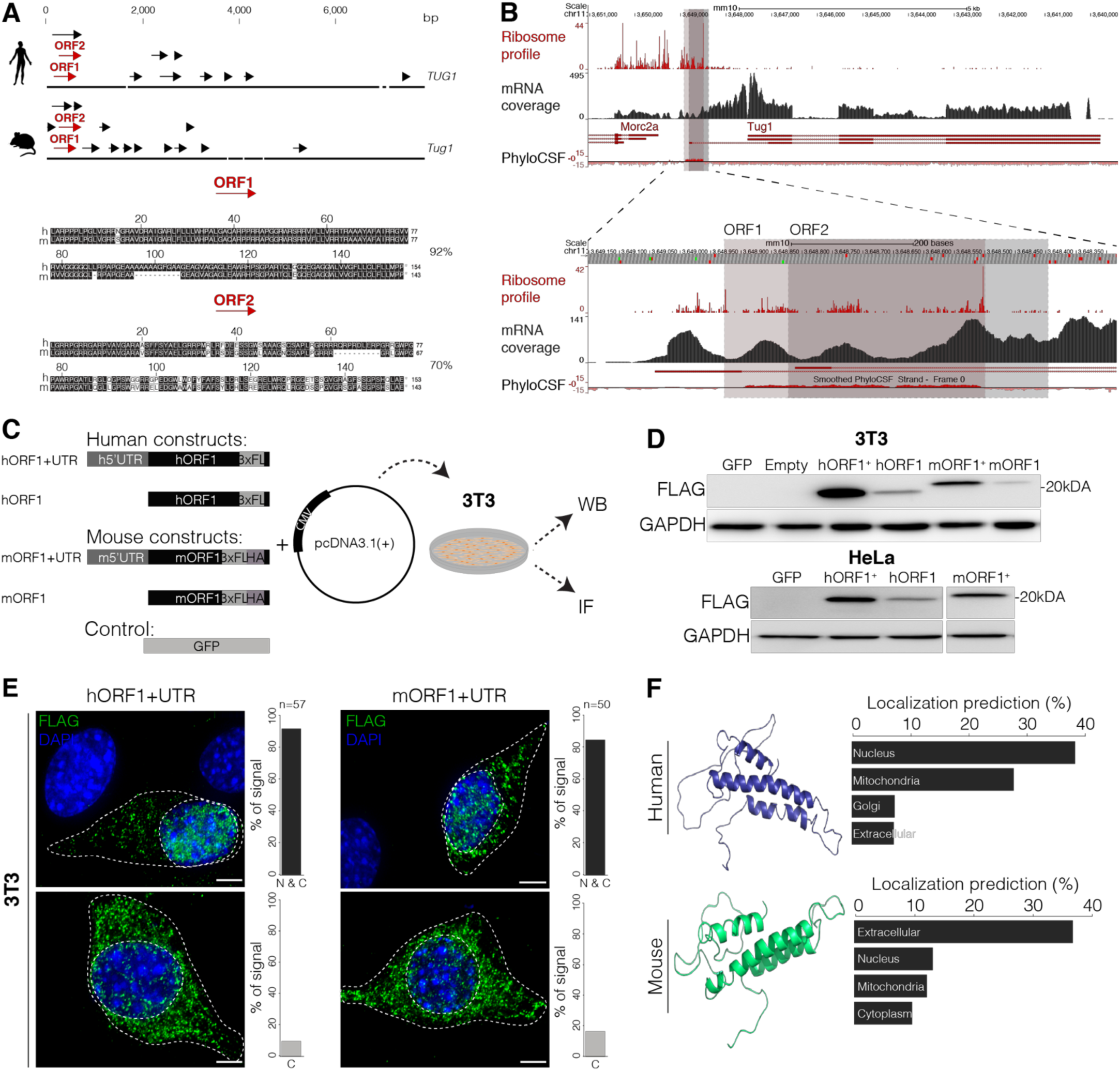
The 5’ region of *Tug1* encodes a conserved peptide. **(A)** Open reading frame (ORF) search in human and mouse *Tug1* reveals multiple ORFs (arrows). ORF1 and ORF2 (depicted as red arrows) indicate two ORFs with greater than 70% amino acid conservation between human and mouse (92% and 70%, respectively). **(B)** *Tug1* mouse genomic locus (mm10) is shown. Ribosome occupancy (Ribosome profile), RNA-seq (mRNA coverage), and evolutionary protein-coding potential (PhyloCSF) across the *Tug1* locus in mouse embryonic fibroblasts (MEFs) is depicted. ORF1 and ORF2 are outlined with red and gray boxes, respectively (top). Tracks surrounding both ORFs are zoomed in for clarity (bottom). **(C)** Scheme of human and mouse ORF1 construct design. A 3xFLAG epitope tag was inserted prior the stop codon of ORF1. Mouse constructs were dual-tagged with both 3xFLAG and HA tags. Expression constructs were designed with (hORF1+UTR, mORF1+UTR) and without (hORF1, mORF1) the 5’ UTR. Constructs and GFP as control were inserted into pcDNA3.1(+). 48 hours post-transfection, 3T3 and HeLa cells were harvested for western blot (WB) (shown in D) or analyzed by immunofluorescence (IF) (shown in E). **(D)** Western blot targeting the 3xFlag (FLAG) in 3T3 and HeLa cells expressing human and mouse constructs, respectively. GAPDH was used as a loading control. **(E)** Maximum intensity projections of 3T3 cells expressing human and mouse constructs. Immunostaining against the Flag tag (green) and DAPI (blue) are shown. Bar plot shows localization analysis of human and mouse TUG1-BOAT. N and C indicates nuclear and cytoplasmic localization, C indicates only cytoplasmic. Scale bar is 5 μm. **(F)** Human and mouse TUG1-BOAT structure (RaptorX) and localization (DeepLoc) prediction.

To further hone in on translated regions of *Tug1*, we analyzed ribosome profiling data (Michel et al., 2014), which identifies regions of RNA bound to ribosomes by high-throughput sequencing, thus indicating actively translating portions of an RNA. We found pronounced ribosomal occupancy across the entire ORF1 sequence with a sharp decrease at its stop codon (Figure 5B) (Ingolia et al., 2009). A similar pattern indicative of active translation of *Tug1* ORF1 is also observed from ribosome profiling in human, mouse, and rat heart tissue (S. van Heesch, personal communication, September 2018). However, ORF2 does not show ribosome occupancy above background level, particularly after the ORF1 stop codon (Figure 5B and Figure S6A). Taken together, these results suggest that the most 5’ region of *TUG1*/*Tug1* contains an ORF that has evolutionary conservation characteristic of protein-coding genes. We designated the putative peptide originating from ORF1 as TUG1-BOAT (*Tug1*-**B**ifunctional **O**RF **a**nd **T**ranscript).

To determine if ORF1 is translated, we first performed *in vitro* translation assays using [35S]-methionine incorporation to detect newly synthesized proteins for three different constructs: (i) the endogenous *TUG1* lncRNA (including the endogenous 5’UTR, ORF1 and a part of the 3’UTR), (ii) a codon optimized ORF1-3xFLAG and (iii) a codon optimized ORF1-mEGFP (Figure S6B). For each construct, we observed a protein product of the expected size, thereby supporting that ORF1 can produce a stable peptide (Figure S6C). We next generated C-terminal epitope tagged human and mouse TUG1-BOAT expression constructs with and without the 5’ leader sequences (Figure 5C). As a negative control, we generated a construct containing GFP in place of the TUG1-BOAT cDNA sequence. We then transfected 3T3 (mouse) and HeLa (human) cells and tested for TUG1-BOAT translation by western blot analysis. We detected peptides of approximately 19 kDa and 21 kDa in both cell lines (Figure 5D), which closely corresponds to the predicted molecular weights of hTUG1-BOAT (18.7 kDa) and mTUG1-BOAT (19 kDa) fusion constructs, respectively. Collectively, these results show that ORF1, with its 5’ UTR and a native non-canonical translational start site, can be translated into TUG1-BOAT in both human and mouse cells.

Having detected a peptide of expected size from human and mouse TUG1-BOAT constructs, we next investigated the peptide’s subcellular localization by immunofluorescence. We observed that human and mouse TUG1-BOAT is distributed throughout the nucleus and cytoplasm in the majority of the cells (>80 % cells, n = 50) (Figure 5E). However, in a subset of cells, TUG1-BOAT was predominantly cytoplasmic (<20 % of cells, n = 50) (Figure 5E). Moreover, we found that TUG1-BOAT showed co-localization with the mitochondria (Figure S6D).

### TUG1-BOAT overexpression compromises mitochondrial membrane potential

We next sought to identify a potential cellular role for TUG1-BOAT. We used protein structure/domain prediction tools to further characterize TUG1-BOAT. Based on predictions, TUG1-BOAT does not represent any known homologs, and the predicted structures are conserved between human and mouse (template modeling score of 0.658). Further investigation of putative functional domains revealed a conserved mitochondrial localization domain (Figure 5F). Based on the predicted mitochondrial localization domain encoded in TUG1-BOAT (Figure 5F), its co-localization with the mitochondria (Figure S6D), and given that oxidative phosphorylation was one of the most affected pathways across multiple *Tug1*^-/-^ tissues (Figure 4C), we hypothesized that TUG1-BOAT may have a role in the mitochondria.

To this end, we first examined mitochondrial membrane potential by using chloromethyl-X-rosamine (CMXR), a lipophilic fluorescent cation that accumulates in the negatively charged interior of mitochondria (Macho et al., 1996). We transfected human and mouse TUG1-BOAT expression constructs with and without the 5’ UTR, as well as a control GFP-containing plasmid and a *Tug1* construct that lacks ORF1 (*Tug1* cDNA ΔmORF1) into 3T3 cells (Figure 6A). Notably, cells with either human or mouse TUG1-BOAT showed a reduction in mitochondrial staining by CMXR (22% and 44% CMXR stained cells, respectively), compared to cells in the same culture not expressing TUG1-BOAT (Figure 6B). In contrast, cells expressing GFP or *Tug1* cDNA ΔmORF1 were positive for CMXR staining in all cells examined, thus indicating that CMXR staining deficiency is induced by the TUG1-BOAT peptide alone, rather than the *Tug*1 RNA.

**Figure 6.**
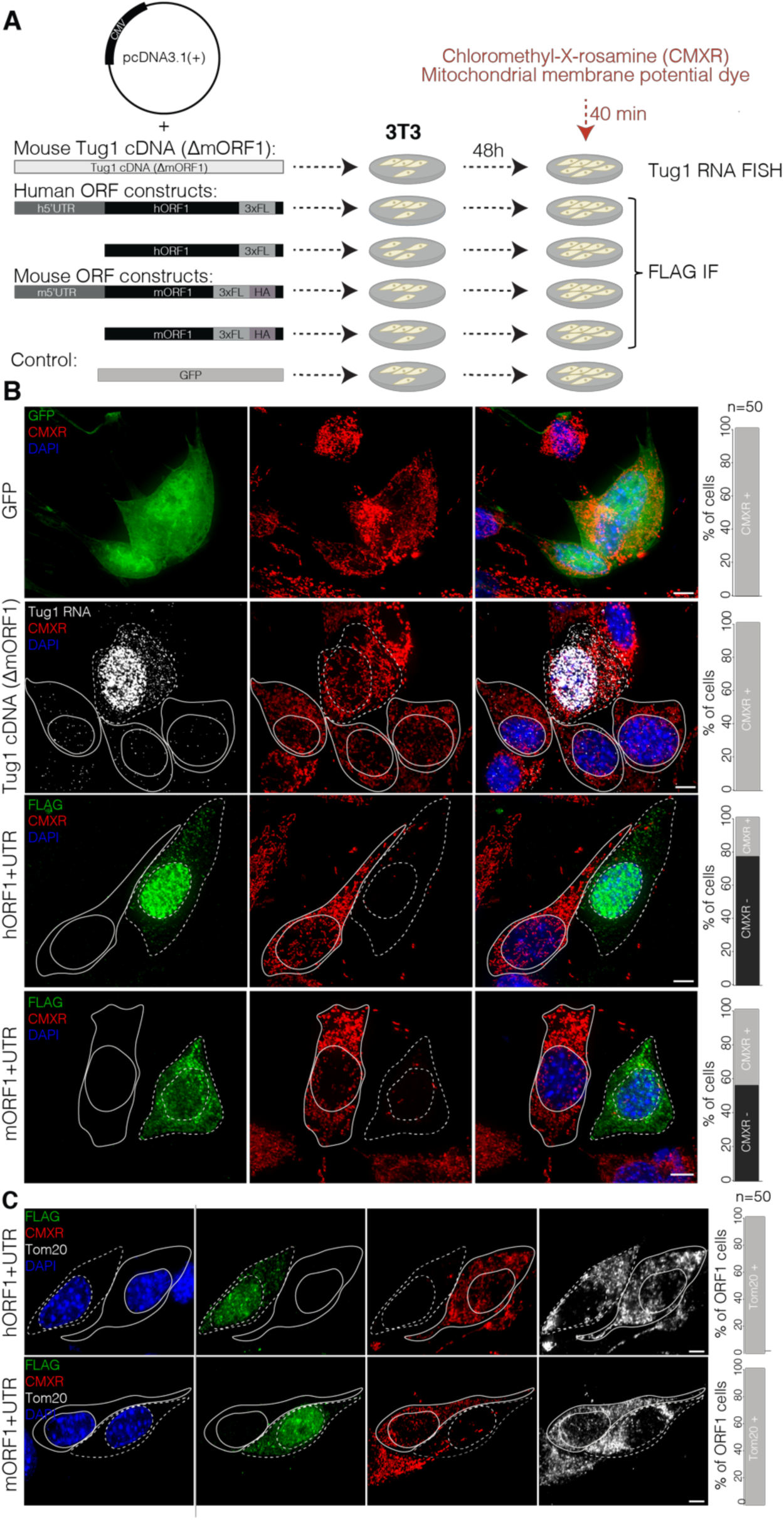
Overexpression of TUG1-BOAT compromises mitochondrial membrane potential. **(A)** Construct and transfection scheme. Human and mouse ORF1, and mouse *Tug1* cDNA lacking the ORF1 region (*Tug1* cDNA ΔmORF1) were inserted into pcDNA3.1(+). Chloromethyl-X-rosamine (CMXR) was added to visualize mitochondria 48 hours post transfection. After staining, cells were fixed and processed for anti-FLAG immunofluorescence (IF) or *Tug1* RNA FISH. **(B)** Maximum intensity projections of z-stacks acquired 48 hours post-transfection of 3T3 cells with indicated plasmids and staining with CMXR. *Tug1* RNA overexpression was monitored by *Tug1* single molecule RNA FISH (gray), TUG1-BOAT by immunostaining against the FLAG tag (green). GFP was used as a control. CMXR is shown in red, DAPI in blue. On the right, quantification of cells positive for GFP, *Tug1* RNA, or TUG1-BOAT and mitochondria by CMXR (n = 50). Scale bar is 5 μm. Maximum intensity projections of z-stacks acquired 48 hours post-transfection of 3T3 cells with the indicated plasmids, stained with CMXR (red) and immunostained against mitochondrial membrane translocase TOM20 (gray). On the right, quantification of cells over-expressing TUG1-BOAT and lacking CMXR staining showing intact mitochondrial membrane assessed by TOM20. Nuclei were stained with DAPI (blue). Scale bar is 5 μm

Since CMXR is commonly used to measure mitochondrial membrane potential, we reasoned that either impaired mitochondrial integrity or impaired redox potential at the mitochondrial membrane could account for the accumulation defect of CMXR in mitochondria upon TUG1-BOAT overexpression. To address these possibilities, we immunostained for TOM20, a redox independent translocase located on the outer mitochondrial membrane (Likić et al., 2005) in cells overexpressing human or mouse TUG1-BOAT. We observed staining for TOM20 in cells without CMXR staining, indicating that the mitochondria were intact (Figure 6C). Collectively, these results provide evidence that human and mouse TUG1-BOAT have conserved roles in mitochondrial function, and that *Tug1* RNA, DNA, and the TUG1-BOAT peptide have distinct roles.

## DISCUSSION

To date, there are a few well-established *in vivo* genetic models of lncRNAs with robust phenotypes and lncRNAs remain understudied, as a class, in this context. This is further complicated by the fact that lncRNA loci can contain multiple regulatory modalities including the DNA, RNA, protein, and the act of transcription. Therefore, when a lncRNA locus presents a robust phenotype, understanding what molecular activities are present at the locus is an important foundation in order to then address how it could potentially mediate an effect. In this study, we characterized in more detail one of our previously published lncRNA knockout mouse models, *Tug1*, and extended our understanding of the function of this locus *in vivo* by defining the molecular properties present at the locus. By implementing multiple *in vivo* genetic strategies, we report a number of key findings: (i) in our mouse model, deletion of the *Tug1* locus leads to completely penetrant male sterility due to late stage spermatogenesis defects, (ii) we find evidence that *Tug1* harbors a *cis*-acting DNA repressive element, and (iii) we find evidence that the *Tug1* RNA regulates a subset of genes in *trans*. Moreover, using biochemical and cell-based assays, we find evidence that (iv) *Tug1* encodes a highly conserved peptide in its 5’ region that appears to have a mitochondrial role. Together, our results point to an essential role for the *Tug1* locus in male fertility, where the locus harbors three distinct regulatory activities.

### LncRNAs in spermatogenesis and male fertility

Infertility is estimated to affect approximately 15% of couples in developed countries, with male infertility contributing up to 50% of cases. Much about the molecular regulation of male germ cell development remains to be understood, especially the contribution of the noncoding genome. Indeed, a number of studies have performed systematic gene expression profiling at defined stages during spermatogenesis and have identified many developmentally regulated lncRNAs, suggesting that lncRNAs may have a wide role during spermatogenesis (Wichman et al., 2017). In our gene ablation knockout for *Tug1*, we observed a male sterility phenotype with complete penetrance. Although four other lncRNAs (*Tslrn1, Tsx, Pldi*, and *Mrlh*) have been found to be important in spermatogenesis, none of them lead to male sterility when disrupted (Anguera et al., 2011; Arun et al., 2012; Heinen et al., 2009; Wichman et al., 2017).

Our stage-specific analysis of *Tug1*^-/-^ sperm identified several key abnormalities. We observed defects of cytoplasm removal during spermiation, causing mechanical strain on the midpiece region of sperm that include the point of attachment with both head and tailpiece. Interestingly, similar aberrant sperm phenotypes associated with defects of cytoplasm removal during spermiation have been described for the protein-coding genes *Spem1* and *Ehd1* in loss-of-function models (Rainey et al., 2010; Zheng et al., 2007). The timing and mechanism by which sperm shed their collective cytoplasm during individualization is highly regulated (O’Donnell et al., 2011; Steinhauer, 2015); however, this process is poorly characterized in mammals. Based on data from earlier studies, we speculate that the sterility of *Tug1*^-/-^ males arises from a combination of oligozoospermia (low sperm count) and teratozoospermia (abnormal morphology) resulting from a failure of spermatids to individualize during spermatogenesis.

### Multiple molecular modalities of the Tug1 locus

Toward understanding the molecular activities present at the *Tug1* locus, we investigated the activity of the DNA, lncRNA, and peptide and found that the *Tug1* locus harbors three distinct regulatory activities. First, several lines of evidence indicate that the *Tug1* locus has a *cis*-acting repressive DNA function. We observed that upon deletion of the *Tug1* locus, genes downstream of *Tug1* were consistently upregulated across multiple tissues, and this effect was also observed in an allele-specific manner – thereby suggesting that the local repressive *cis*-effect is mediated by DNA regulatory elements within the *Tug1* locus. Indeed, DNA regulatory elements have been found overlapping gene bodies, whether protein coding or noncoding. Contrary to enhancers, only a handful of repressor elements and silencers have been identified and characterized to date in mammalian genomes (Li and Arnosti, 2011; Li et al., 2014; Qi et al., 2015; Tan et al., 2010). In keeping with the idea of a repressive role for the DNA at the *Tug1* locus, it is notable that in a recent study which systematically tested for enhancer activity across lncRNA loci, enhancer activity was detected at all lncRNA loci examined except for *Tug1*, where only the promoter showed activity (Groff et al., 2018). Further defining the precise DNA repressive elements as well as their mechanism will be of interest to understand the regulatory abilities of the DNA elements within the *Tug1* locus.

Second, our study finds regulatory activity for the *Tug1* lncRNA and thus extends previous findings for the role of *Tug1* RNA on gene expression (Long et al., 2016). Using compound transgenic mouse models, we found that a subset of genes found dysregulated in *Tug1*^-/-^ testes, could be reciprocally regulated by ectopic expression of *Tug1* RNA, even at low levels. While our transgene was expressed at lower levels than wild type *Tug1* RNA, other lncRNAs such as *Hottip* and *Xist* have been shown to exert a biological activity at relatively low copy numbers (Sunwoo et al., 2015; Wang et al., 2011). In support of a *trans*-acting role for *Tug1* RNA, a previous study found that *Tug1* RNA can regulate the levels of *Ppargc1a* mRNA in a reciprocal manner in cultured podocytes (Long et al., 2016). In our RNA-seq dataset of eight different tissues we did not find significant dysregulation for *Ppargc1a*, but it is important to note that our RNA-seq dataset does not include the kidney. Expressing *Tug1* RNA at higher levels could uncover a more widespread *trans* role for *Tug1* to regulate gene expression and is of interest for future work.

Finally, we demonstrate that *Tug1* encodes an evolutionarily conserved peptide in the 5’ region that, when overexpressed, impacts mitochondrial membrane potential. TUG1-BOAT has high positive charge (net charge ∼ +16.5), thus we speculate that the high accumulation of such a positively charged peptide at the mitochondria in the overexpression experiments could lead to depolarization of the mitochondrial membrane. A number of recent studies have identified peptides at candidate lncRNA loci, thereby highlighting the complexity of these loci (Anderson et al., 2015; Chng et al., 2013; Nelson et al., 2016; Stein et al., 2018). In one recent example, the candidate lncRNA *LINC00661* was shown to encode a conserved peptide that has a role in mitochondrial function (Stein et al., 2018). In support of a mitochondrial role for the *Tug1* locus, a previous report found that overexpression of a *Tug1* isoform impacted mitochondrial bioenergetics in cultured podocytes from a murine diabetic nephropathy model (Long et al., 2016).

### The Tug1 locus has multiple regulatory modalities with potential function in spermatogenesis

Collectively, our study identifies that deletion of the *Tug1* locus results in a completely penetrant male sterility phenotype, and that the *Tug1* locus contains three unique regulatory modalities: the DNA repressive element, the *Tug1* lncRNA transcript, and the peptide (TUG1-BOAT). As such, our findings pose an intriguing possibility that these features could individually or in combination mediate the observed fertility defect in *Tug1* knockout mice. While our study does not resolve this outstanding question, there is evidence to support a role for each modality for further investigation. First, there is some evidence that the cohort of genes downstream of the *Tug1* locus that are transcriptionally upregulated when *Tug1* is deleted have a role in male fertility. Two of the six dysregulated genes in *Tug1*^-/-^ testes, *Smtn* and *Pla2g3*, have loss-of-function mutations that have male fertility and sperm maturation defects (Niessen et al., 2005; Sato et al., 2010). However, links between these genes and fertility in an overexpression context in an animal model have not been reported. In addition, the effect on fertility for the other four genes (*Gm11946, Rnf185, Selm*, and *8430429K09Rik*) that were found upregulated upon *Tug1* deletion have not been reported in either loss-of-function or overexpression contexts. Second, in support of a potential role for the *Tug1* lncRNA in male fertility, it is notable that *Selenop*, a gene found up-regulated in *Tug1*^-/-^ testes and was reciprocally regulated in the *Tug1*^*rescue*^ testes, has a known role in male fertility in the loss-of-function context (Hill et al., 2003). Yet, the role of *Selenop* on male fertility in an overexpression context has yet to be reported. Moreover, given that our *Tug1*^rescue^ mice did not restore fertility, this finding may indicate that it is not due to the lncRNA; however, the lack of a rescue may also be explained by the low levels of *Tug1* expression from the transgene. Finally, in our TUG1-BOAT experiments we observed altered mitochondrial membrane potential, which has also been observed in male sterility (Wang et al., 2003). Thus, the TUG1-BOAT peptide may also play a role in male fertility.

## CONCLUSIONS

Our findings reveal an essential role for *Tug1* in male fertility, providing evidence that *Tug1* knockout male mice are sterile with complete penetrance due to a low sperm count and abnormal sperm morphology. Moreover, we show that the *Tug1* locus harbors three distinct regulatory activities that could account for the fertility defect, including (i) a *cis* DNA repressor that regulates many neighboring genes, (ii) a lncRNA that can regulate genes by a *trans*-based function, and (iii) an evolutionary conserved peptide that when overexpressed impacts mitochondrial membrane potential. Thus, our study provides a roadmap for future studies to investigate the individual and/or combined contributions of *Tug1* DNA, RNA, and/or peptide to the male fertility defect, as well as in additional diseases in which *Tug1* is altered.

## METHODS

### Mice and ethics statement

Mice used in these studies were maintained in a pathogen-specific free facility under the care and supervision of Harvard University’s Institutional Animal Care Committee. *Tug1*^*tm1.1Vlcg*^ knockout mice have been described previously (Goff et al., 2015; Sauvageau et al., 2013). To remove the *loxP*-flanked neomycin resistance gene included in the targeting construct, we crossed *Tug* ^*tm1.1Vlcg*^ mice to *C57BL6/J* mice and then to a cre-recombinase strain (*B6.C-Tg*^*(CMV-cre)1Cgn/J*^, The Jackson Laboratory, 006054). Mice free of both the neomycin-resistance and cre-recombinase genes were selected for colony expansion and subsequently backcrossed to *C57BL/6J* mice. The *Tug1* knockout allele was maintained by heterozygous breeding, and mutant mice were identified by genotyping for loss of the *Tug1* allele and gain of the *lacZ* cassette (Transnetyx, Inc.).

For allele specific gene expression analyses, we generated *Tug1*^BL6-KO/Cast-WT^ mice by crossing inbred *Mus castaneus* (Cast/EiJ) males (The Jackson Laboratory, 000928) with inbred heterozygote *Tug1* females. The F1 hybrid male progeny (three wild type *Tug1*^BL6-WT/Cast-WT^ and four with a maternal *Tug1* knockout allele *Tug1*^BL6-KO/Cast-WT^) were used for allele-specific expression studies.

To generate an inducible *Tug1*-overexpression mouse, tg(*Tug1*), we cloned *Tug1* cDNA (see Sequences and Primers below) into a Tet-On vector (pTRE2). Full length *Tug1* (Ensembl id: ENSMUST00000153313.2) was amplified from Riken cDNA clone E330021M17 (Source Bioscience) using specific primers containing MluI and EcoRV restriction sites (see Sequences and Primers below). After gel purification, we subcloned the amplicon using the MluI and EcoRV restriction sites into a modified Tet-On pTRE2pur vector (Clontech 631013) in which the bGlobin-intron was removed. We verified the absence of mutations from the cloned *Tug1* cDNA by sequencing (see Sequences and Primers below). We injected this cassette into the pronucleus of C57BL/6J zygotes (Beth Israel Deaconess Medical Center Transgenic Core). Two male founder mice containing the tg(*Tug1*) cassette were identified by genotyping for the pTRE allele and individually mated to female C57BL/6J mice (Jackson Laboratory, 000664) to expand the colonies. Next, we generated quadruple allele transgenic mice to test the functionality of the *Tug1* RNA by the following strategy. We mated tg(*Tug1*) males to *Tug1*^*tm1.1Vlcg*^ females and identified male progeny that were *Tug1*^+/-^; tg(*Tug1*). These mice were then mated to female rtTA mice (B6N.FVB(Cg)-Tg(CAG-rtTA3)4288Slowe/J mice (Jackson Laboratory, 016532)) and we identified male progeny that were *Tug1*^+/-^; tg(*Tug1*), *rtTA*. Finally, we mated male *Tug1*^+/-^; tg(*Tug1*), *rtTA* mice to *Tug1*^+/-^ females, and at the plug date, females were put on 625 mg/kg doxycycline-containing food (Envigo, TD.01306). We genotyped progeny from the above matings (Transnetyx, Inc) and identified male progeny that were *Tug1*^-/-^; tg(*Tug1*), *rtTA*, and maintained these mice on the doxycycline diet until the experimental end point.

### Cell Lines and Cell Culture

We derived primary wild type and *Tug1*^*-/-*^ mouse embryonic fibroblasts (MEFs) from E14.5 littermates from timed *Tug1*^*+/-*^ intercrosses as described (Xu, 2005). We maintained MEFs as primary cultures in DMEM, 15% FBS, pen/strep, L-glutamine and non-essential amino acids. We genotyped MEFs derived from each embryo and used only male *Tug1*^*-/-*^ and wild type littermate MEFs at passage 2 for all experiments.

3T3 (ATCC, CRL-1658™), HeLa (ATCC, CRM-CCL-2), and BJ (ATCC, CRL-2522™) cell lines were purchased from ATCC and cultured as recommended.

### Whole Mount *In Situ* Hybridization

We generated an antisense riboprobe against *Tug1* (see Sequences and Primers below) from plasmids containing full length *Tug1* cDNA (Ensembl id: ENSMUST00000153313.2) and performed *in situ* hybridization on a minimum of three *C57BL6/J* embryos per embryonic stage. For whole-mount staining, we fixed embryos in 4% paraformaldehyde for 18 hours at 4°C, followed by three washes for 10 minutes each in PBS. We then dehydrated embryos through a graded series of 25%, 50%, 75% methanol / 0.85% NaCl incubations and then finally stored embryos in 100% methanol at −20°C before *in situ* hybridization. We then rehydrated embryos through a graded series of 75%, 50%, 25%, methanol/ 0.85% NaCl incubations and washed in 2X PBS with 0.1% Tween-20 (PBST). Embryos were treated with 10mg/mL proteinase K in PBST for 10 minutes (E8.0, E9.5) or 30 minutes (E10.5, E11.5 and E12.5). Samples were fixed again in 4% paraformaldehyde/0.2% glutaraldehyde in PBST for 20 minutes at room temperature and washed in 2X PBST. We then incubated samples in pre-hybridization solution for 1 hour at 68°C and then incubated samples in 500 ng/mL of *Tug1* antisense or sense riboprobe at 68°C for 16 hours. Post-hybridization, samples were washed in stringency washes and incubated in 100 μg/mL RNaseA at 37°C for 1 hour. Samples were washed in 1X maleic acid buffer with 0.1% Tween-20 (MBST) and then incubated in Roche Blocking Reagent (Roche, #1096176) with 10% heat inactivated sheep serum (Sigma, S2263) for 4 hours at room temperature. We used an anti-digoxigenin antibody (Roche, 11093274910) at 1:5000 and incubated the samples for 18 hours at 4°C. Samples were washed 8 times with MBST for 15 min, 5 times in MBST for 1 hour, and then once in MBST for 16 hours at 4°C. To develop, samples were incubated in 3X NTMT (100 mM NaCl, 100 mM Tris-HCl (pH 9.5), 50 mM MgCl2, 0.1% Tween-20, 2 mM levamisole). The *in situ* hybridization signal was developed by adding BM Purple (Roche, 11442074001) for 4, 6, 8, and 12 hours. After the colorimetric development, samples were fixed in 4% paraformaldehyde and cleared through a graded series of glycerol/1X PBS and stored in 80% glycerol. Finally, we imaged embryos on a Leica M216FA stereomicroscope (Leica Microsystems) equipped with a DFC300 FX digital imaging camera.

### *Tug1* Single Molecule RNA FISH

We performed *Tug1* single molecule RNA FISH as described previously (Raj et al., 2008). Briefly, 48 oligonucleotides labeled with Quasar 570 and Quasar 670 tiled across human/mouse *Tug1* transcripts were designed with LGC Biosearch Technologies’ Stellaris probe designer (Stellaris® Probe Designer version 4.2) and manufactured by LGC Biosearch Technologies.

Human foreskin fibroblasts (ATCC® CRL-2522™) and mouse 3T3 fibroblasts (ATCC, CRL-1658™) were seeded on glass coverslips previously coated with poly-L-lysine (10 μg/mL) diluted in PBS. Prior to hybridization, coverslips were washed twice with PBS, fixed with 3.7% formaldehyde in PBS for 10 minutes at room temperature, and washed twice more with PBS. Coverslips were immersed in ice-cold 70% EtOH and incubated at 4°C for a minimum of 1 hour. We then washed the coverslips with 2 mL of Wash buffer A (LGC Biosearch Technologies) at room temperature for 5 minutes. Next, we hybridized cells with 80 μL hybridization buffer (LGC Biosearch Technologies) containing *Tug1* probes (1:100) overnight at 37°C in a humid chamber. The following day, we washed cells with 1 mL of wash buffer A for 30 minutes at 37°C, followed by another wash with wash buffer A containing Hoechst DNA stain (1:1000, Thermo Fisher Scientific) for 30 minutes at 37°C. Coverslips were washed with 1 mL of wash buffer B (LGC Biosearch Technologies) for 5 minutes at room temperature, mounted with ProlongGold (Life Technologies) on glass slides and left to curate overnight at 4°C before proceeding to image acquisition (see below).

### Sperm Counts and Morphology

*Tug1*^*-/-*^ (n=8) and wild type (n=9) males between 8 and 41 weeks of age were sacrificed and weighed. We then dissected the entire male reproductive tract in phosphate buffered saline (PBS). One testis was removed, weighed and fixed in 4% paraformaldehyde (PFA) for histology (see below). Sperm were collected from one cauda epididymis by bisecting and suspending the tissue in a solution of Biggers-Whitten-Whittingham (BWW) sperm media at 37°C. After a 15-minute incubation, we used the collected sperm solutions to analyze sperm morphology and counts.

We characterized sperm morphology by fixing sperm in 2% PFA in PBS, mounting 20 μL of suspended sperm in Fluoromount-G media (Southern Biotech) on superfrost glass slides (Thermo Fisher Scientific) and scanning each slide in a linear transect, recording the morphology as normal or abnormal for each sperm cell encountered (between 30 to 120 sperm). When abnormal, we also recorded the type of morphological defects: headless, head angle aberrant, head bent back to midpiece, debris on head, debris on hook, head misshapen, midpiece curled, midpiece kinked, midpiece stripped, debris on midpiece, tailless, tail curled, tail kinked, tail broken, or multiple cells annealed together.

Sperm counts for each *Tug1*^*-/-*^ (*n* = 7) and wild type (*n* = 9) mice were determined using a Countess Automated Cell Counter according to manufacturer’s protocol (Life Technologies, Carlsbad, CA). For the *Tug1*^rescue^ experiment, sperm counts for control (WT and *Tug1*^+/-^) (*n* = 2), *Tug1*^-/-^ (n = 2), and *Tug1*^-/-^; tg(*Tug*1); *rtTA* mice (*n* = 3) was determined by manual counts using a hemocytometer. For all analyses, statistical comparisons between *Tug1*^*-/-*^ and wild type was performed using the two-tailed Wilcoxon rank sum tests with an a = 0.05. Results for testes, sperm counts and morphological parameters are presented in Extended Data Table 1. All statistical comparisons of *Tug1*^*-/-*^ versus wild type for relative testis size, sperm morphology and sperm counts were performed using R (Wilcoxon rank-sum test, and principal component analysis (PCA)).

### *lacZ* and Histological Staining of Male Reproductive Tissues

Expression of the knock-in *lacZ* reporter and histological staining for morphological analysis of male reproductive tissues was conducted on testes and epididymides from *Tug1*^*-/-*^ (n = 2) and wild type (n = 2) mice. We fixed testis and epididymis in 4% paraformaldehyde in PBS overnight at 4°C and washed tissues three times in PBS. For *lacZ* staining, we rinsed *Tug1*^*+/-*^ and wild type tissues three times at room temperature in PBS with 2 mM MgCl2, 0.01% deoxycholic acid, and 0.02% NP-40. We performed X-gal staining by incubating the tissues for up to 16 hours at 37°C in the same buffer supplemented with 5 mM potassium ferrocyanide and 1 mg/mL X-gal. The staining reaction was stopped by washing three times in PBS at room temperature, followed by 2 hours post-fixation in 4% paraformaldehyde at 4°C.

We then embedded organs in paraffin, sectioned the organs at 6 μm thickness, and then mounted sectioned samples onto glass microscope slides. Testis sections were additionally stained with Mayer’s Hematoxylin, Periodic Acid and Schiff’s Reagent (VWR, 470302-348), and epididymis sections were stained with eosin (VWR, 95057-848). Images were collected using a Zeiss AxioImager.A1 upright microscope or on an Axio Scan Z.1 (Zeiss).

### RNA Isolation and RNA-Seq Library Preparation

We isolated total RNA from mouse tissues, mouse embryonic fibroblasts (MEFs), and blood cells using TRIzol (Life Technology, Carlsbad, CA) by chloroform extraction followed by spin-column purification (RNeasy mini or micro kit, Qiagen) according to the manufacturer’s instructions. RNA concentration and purity were determined using a Nanodrop. We assessed RNA integrity on a Bioanalyzer (Agilent) using the RNA 6000 chip. High quality RNA samples (RNA Integrity Number ≥ 8) were used for library preparation. We then constructed mRNA-seq libraries using the TruSeq RNA Sample Preparation Kit (Illumina) as previously described (Sun et al., 2013). The libraries were prepared using 500 ng of total RNA as input and a 10-cycle PCR enrichment to minimize PCR artifacts. Prior to sequencing, we ran libraries on a Bioanalyzer DNA7500 chip to assess purity, fragment size, and concentration. Libraries free of adapter dimers and with a peak region area (220-500 bp) ≥ 80% of the total area were sequenced. We then sequenced individually barcoded samples in pools of 6, each pool including *Tug1* mutant and wild type samples, on the Illumina HiSeq platform using the rapid-full flow cell with the 101 bp paired-end reads sequencing protocol (Bauer Core, Harvard University FAS Center for System Biology).

### RNA-seq and Gene Set Enrichment Analyses

We mapped sequencing reads to the reference mouse genome (GRCm38) by STAR (Dobin et al., 2013) with the gene annotation obtained from GENCODE (vM16). We counted uniquely-mapped reads for genes by featureCounts (Liao et al., 2014) and calculated TPM (Transcripts Per Million) for genes to quantify gene expression level after normalization of sequencing depth and gene length. Clustering of gene expression was done with Ward’s method using Jensen-Shannon divergence between tissues as distance metric. The R package, Philentropy was used for calculation (Drost, 2018),

We identified differentially-expressed genes by comparing the read counts of biological replicates between the groups using the generalized linear model. Statistical significance was calculated with the assumption of the negative binomial distribution of the read counts and the empirical estimation of variance by using the R packages DESeq2 (Love et al., 2014) and fdrtool (Strimmer, 2008). The genes were filtered if their read counts were less than three in every biological replicate. The genes were called significant if their FDR-adjusted p-values were smaller than 0.05.

We performed Gene Set Enrichment Analysis (GSEA) to evaluate the enrichment of the gene sets available from MSigDB (Subramanian et al., 2005) after mapping genes to gene sets by gene symbols. The statistical significance of a gene set was calculated with the test statistics of individual genes computed by DESeq2. If the FDR-adjusted p-value is less than 0.1, the term was called as significant. We did this calculation using the R package, CAMERA (Wu and Smyth, 2012).

### Allele-Specific Gene Expression Analysis

We performed allele-specific expression analysis as previously described (Perez et al., 2015). For mouse testes samples, we created a *C57BL/6J, Cast/EiJ* diploid genome by incorporating single nucleotide polymorphisms and indels (obtained from the Mouse Genome Project: ftp://ftp-mouse.sanger.ac.uk/REL-1303-SNPs_Indels-GRCm38) from both strains into the *M. musculus* GRCm38 reference genome sequence. We created a transcriptome annotation set as follows. The gencode.vM2.annotation GTF file was downloaded and Mt_rRNA, Mt_tRNA, miRNA, rRNA, snRNA, snoRNA, Mt_tRNA_pseudogene, tRNA_pseudogene, snoRNA_pseudogene, snRNA_pseudogene, scRNA_pseudogene, rRNA_pseudogene, miRNA_pseudogene were removed (not enriched in our RNA-seq libraries). To create an extensive set of transcripts, we added to the gencode.vM2.annotation all transcripts from the UCSC knownGene mm10 annotation file, which are not represented in the gencode.vM2.annotation set. We also added all functional RNAs from the Functional RNA database (fRNAdb) (Mituyama et al., 2009), which did not intersect with any of the previously incorporated transcripts. From this, we then used the UCSC liftOver utility to generate a *C57BL/6J*, Cast/EiJ diploid transcriptome set.

Each RNA-seq library was first subjected to quality and adapter trimming using the Trim Galore utility (http://www.bioinformatics.babraham.ac.uk/projects/trim_galore) with stringency level 3. We then mapped each of the *C57BL/6J∷Cast/EiJ* hybrid RNA-seq libraries to the *C57BL*/*6J* and *Cast*/*EiJ* diploid genome and transcriptome splice junctions using STAR RNA-seq aligner (Dobin et al., 2013), allowing a maximum of 3 mismatches. The data were mapped twice, where after the first mapping step we incorporated valid splice junctions that were reported by STAR to exist in the RNA-seq data. We then transformed the genomic alignments to transcriptomic alignments. Following that, we estimated the expression levels with their respective uncertainties for each transcript in our *C57BL/6J* and *Cast/EiJ* diploid transcriptome using MMSEQ (Turro et al., 2011). The posterior FPKM samples were transformed to TPM units with a minimum expression TPM cutoff set to 0.01. In any RNA-seq sample, any transcript for which its MMSEQ posterior median TPM was lower than 0.01 was set to 0.01 (used as the minimal measurable expression level).

We adopted the approach of Turro et al. for combining lowly identifiable transcripts based on the posterior correlation of their expression level estimates, tailored for a diploid transcriptome case (Turro et al., 2014). In this approach, for any given RNA-seq sample we compute the Pearson correlation coefficient of the posterior TPM samples of any pair of transcripts from the same locus and the same allele. Subsequently, if the mean Pearson correlation coefficient across all RNA-seq samples for a pair of transcripts in both alleles is lower than a defined cutoff (which we empirically set to −0.25), each of these pairs is combined into a single transcript. This process continues iteratively until no pair of transcripts (or pairs of already combined transcripts) can be further combined. This consistency between the alleles in the combining process ensures that the resulting combined transcripts are identical for the two alleles and can therefore be tested for allelically biased expression.

### Amplification of Full Length *Tug1*

We amplified the full length *Tug1* isoform lacking the 5’ region (Ensembl id: ENSMUST00000153313.2) from Riken cDNA clone E330021M17 (Source Bioscience) using specific primers containing MluI and EcoRV restriction sites (see Sequences and Primers below). After gel purification, the amplicon was sub-cloned, using the MluI and EcoRV restriction sites, into a modified Tet-On pTRE2pur vector (Clontech, 631013) in which the bGlobin-intron was removed. We verified the absence of mutations from the cloned *Tug1* cDNA by sequencing using primers listed below. The plasmid was used also for sub-cloning *Tug1* into pcDNA3.1(+) (see below).

### ORF Search and TUG1-BOAT Structure and Subcellular Localization Prediction

We analyzed human and mouse *Tug1* cDNA sequences with CLC Genomics Workbench (Qiagen) for open reading frames (ORFs), allowing both canonical and non-canonical start codons (AUG, CUG and UUG). After, sequences with annotated ORFs were aligned using MUSCLE alignment. All further sequence and amino acid alignments were performed with CLC Genomics Workbench.

We predicted secondary and tertiary structure of TUG1-BOAT using RaptorX (Källberg et al., 2012; Peng and Xu, 2011), based on the *Tug1* ORF1 amino acid sequence. RaptorX was chosen for structure prediction due to its ability to predict structures of proteins without known homologs. The resulting PDB files of the predicted structures were visualized using PyMOL. Subcellular localization of human and mouse TUG1-BOAT was predicted with DeepLoc-1.0 (Armenteros et al., 2017).

### Generation of Human and Mouse TUG1-BOAT Overexpression Constructs

We generated a synthesized construct for human *Tug1* ORF1 that contained an in-frame 3xFLAG epitope tag prior to the stop codon, with and without the 5’ leader sequence (GeneWiz). We also synthesized a construct containing mouse ORF1 with an HA tag after the 3xFLAG before the stop codon, with and without the 5’ leader sequence (GeneWiz).

We amplified the *Tug1* cDNA sequence with primers (see Sequences and Primers below) having KpnI and NotI restriction enzyme overhangs from the pTRE2-*Tug1* vector plasmid using Q5 polymerase (Roche) and under following conditions: 96°C for 2 minutes, 35 cycles of (96°C for 30 seconds, 65°C for 30 seconds, 72°C for 4 minutes), 72°C for 4 minutes, and gel purified the amplicon. We digested the inserts and pcDNA3.1(+) plasmid with proper restriction enzymes according to manufacturer’s instructions. After digestion, the plasmid was dephosphorylated using alkaline phosphatase. We then ligated the plasmid and inserts using T4 ligase (NEB) in a 1:3 ratio respectively, followed by bacterial transformation, culture growth, and plasmid isolation (Qiagen Mini-Prep Kit).

### Transfection of TUG1-BOAT Constructs

We seeded 3T3 and HeLa cells in 10 cm plates containing poly-L-lysine coated 18 mm glass cover slips. Next, we transfected the cells with 14 μg of plasmid (pcDNA3.1(+) containing each of the inserts) using Lipofectamine™ 3000 Transfection Reagent (Thermo Fisher Scientific) per manufacturer’s recommendations. 48 hours post transfection, cell pellets were harvested for protein extraction (see below) and coverslips were processed for RNA FISH and/or immunofluorescence (see below).

### Protein Extraction and Western Blot

We resuspended 3T3 and HeLa cell pellets in RIPA Lysis and Extraction Buffer 48 hours post transfection (Thermo Fisher Scientific). Total protein was quantified with Pierce™ BCA® Protein Assay Kit (Thermo Fisher Scientific). We then separated a total of 20-25 μg of denatured protein on a 12.5% SDS polyacrylamide gel for 100 minutes at 120V. We transferred proteins to an Immobilon-PSQ PVDF membrane (Sigma-Aldrich, ISEQ00010) at 400 mA for 75 minutes. After blocking in 5% dried milk in TBST, the membrane was incubated with properly diluted primary antibody (M2 Monoclonal ANTI-FLAG 1:1000, F1804, Sigma; Monoclonal GAPDH 1:5000, 2118S, CST) in 5% dried milk/TBST overnight at 4°C. The next day, we washed the membrane three times for 5 minutes each in TBST (0.5% Tween-20). We then incubated the membrane with Horse Radish Peroxidase-conjugated secondary antibody (Anti-mouse 1:15,000, A9044, Sigma; Anti-rabbit 1:10,000, 711035152, Jackson Immunoresearch), diluted in 5% dried milk/TBST for 1 hour at room temperature. Following three 5 minute washes in TBST, SuperSignal™ West Pico PLUS chemiluminescent substrate (Thermo Scientific, 34580) was added and chemiluminescence was detected using ImageQuant™ LAS 4000 imager.

### Mouse TUG1-BOAT Localization by Immunofluorescence

We plated HeLa and 3T3 cells on poly-L-Lysine coated coverslips. 48 hours post transfection, we rinsed coverslips twice with PBS and fixed cells with 3.7 % formaldehyde in PBS for 10 minutes at room temperature. After 2 washes with PBS, we permeabilized cells with PBT (PBS, 0.1% Tween-20) for 15 minutes at room temperature. Next, we blocked coverslips with 5% BSA in PBT for 1 hour at room temperature and then incubated coverslips with properly diluted primary antibody (mouse M2 monoclonal ANTI FLAG, 1:800, F1804, Sigma; rabbit polyclonal Tom20, 1:800, FL-145, Santa Cruz) in 5 % BSA in PBT for 3 hours at 37°C in a humid chamber. Coverslips were washed three times for 5 minutes each with PBT and incubated with diluted secondary antibody (anti-mouse labelled with Alexa Fluor 488, 1:800, ab150113, Abcam; anti-rabbit labelled with Alexa Fluor 647, 1:800, 4414S, CST) in 5% BSA in PBT for 1 hour at room temperature. Cells were then washed twice for 5 minutes with PBS, once for 20 minutes with PBS containing Hoechst DNA stain (1:1000, Thermo Fisher Scientific), rinsed in PBS, and then mounted on glass slides with ProLong Gold (Thermo Fisher Scientific).

### Mitochondrial Staining with MitoTracker® Red Chloromethyl-X-rosamine

We plated cells on poly-L-lysisne coated coverslips and transfected as described in the previous sections. 48 hours post transfection, cells were incubated with 200 nM MitoTracker® Red Chloromethyl-X-rosamine (Thermo Fischer Scientific, M7512) in 1 mL FBS-free growth media for 40 minutes. We then washed cells twice with PBS, fixed with 3.7% formaldehyde for 10 minutes at room temperature, and processed for immunofluorescence and/or RNA FISH (as described previously).

### *In vitro* Translation of Human *TUG1*

Synthetic gene constructs were produced by Genewiz (constructs available upon request) and designed to capture (i) a selection of the endogenous human *TUG1* lncRNA, which includes the predicted ORF1 with a CUG translation initiation site, the 5’ UTR and 321 nucleotides of the 3’ UTR (chr22:30,969261-chr22:30,970,140), (ii) a codon-optimized sequence for the human *TUG1* translated ORF1 with a 3xFLAG inserted before the termination codon, and (iii) a codon-optimized human *TUG1* translated ORF1 with an mEGFP inserted before the termination codon. The sequence of the translated human *TUG1* lncRNA transcript includes an alternative exon 1 transcriptional start site at chr22:30,969,261 (hg38) obtained from a combination of *de novo* transcriptome assembly publicly available CAGE data. *TUG1* constructs were transcribed and translated *in vitro* from 0.5 μg linearized plasmid DNA using the TnT® Coupled Wheat Germ Extract system (Promega, Mannheim, Germany), in the presence of 10 mCi/mL [35S]-methionine (Hartmann Analytic, Braunschweig, Germany), according to manufacturer’s instructions. 5 μL of lysate was denatured for 2 minutes at 85 °C in 9.6 μL Novex Tricine SDS Sample Buffer (2X) (Thermo Fisher Scientific) and 1.4 μL DTT (500 mM). Proteins were separated on 16% Tricine gels (Invitrogen) for 1 hour at 50 V followed by 3.5 hours at 100 V and blotted on PVDF-membranes (Immobilon-PSQ Membrane, Merck Millipore). Incorporation of [35S]-methionine into newly synthesized proteins enabled the detection of translation products by phosphor imaging (exposure time of 1 day).

### Human TUG1-BOAT Localization by Immunofluorescence

HeLa cells were grown on glass slides for 24 hours and transfected with 3xFLAG-tagged codon optimized human TUG1 ORF1 plasmid using Lipofectamine 2000 reagent for 24 hours. We fixed cells with 4% paraformaldehyde for 10 minutes at room temperature and washed cells three times with ice-cold PBS. The cells were permeabilized and blocked for 1 hour at room temperature using 2.5% bovine albumin serum, 10% anti-goat serum and 0.1% Triton X and washed again. Expressed TUG1 protein was stained for 1 hour at room temperature using a monoclonal anti-FLAG mouse antibody (1:500, F1804, Sigma Aldrich) and co-stained with organelle markers for mitochondria (1:1000, rabbit ATPIF1 #13268, Cell Signaling Technology; Danvers, MA, USA). Afterwards, we washed the slide and incubated with fluorescently-labeled secondary antibodies (1:500, Alexa Fluor 488 anti-rabbit & Alexa Fluor 594 anti-mouse (Invitrogen, Carlsbad, CA, USA) for 30 minutes at room temperature. Cells were washed again, stained with 4-6-diamidino-2-phenylindole (NucBlue™ Fixed Cell ReadyProbes™ Reagent, R37606, Thermo Fisher) for 5 minutes at room temperature and mounted onto glass slides using ProLong™ Gold antifade reagent (Molecular Probes; Invitrogen™). Images were visualized using a LEICA SP8 confocal microscope using a 63x objective. Image analysis was performed using Leica confocal software Las X (v3.5.2) and ImageJ (v1.52a) (Schneider et al., 2012).

### Microscopy and Image Analysis

We acquired z-stacks (200 nm z-step) capturing entire cell volume for single molecule RNA FISH, single molecule RNA FISH/CMXR staining, 3xFLAG tag immunofluorescence/CMXR staining and/or Tom20 immunofluorescence with a GE wide-field DeltaVision Elite microscope with an Olympus UPlanSApo 100x/1.40-NA Oil Objective lens and a PCO Edge sCMOS camera using corresponding filters. 3D stacks were deconvolved using the built-in DeltaVision SoftWoRx Imaging software. Maximum intensity projections of each image were subjected for quantification using Fiji.

### Fluorescence Activated Cell Sorting (FACS)

Age- and sex-matched adult mice were used in all flow cytometry experiments. We obtained peripheral blood by cardiac puncture and collected blood into a 1.5 mL Eppendorf tube containing 4% citrate solution. Next, we added the blood-citrate mixture to 3 mL of 2% dextran/1X PBS solution and incubated for 30 minutes at 37°C. The upper layer was transferred to a new 5 mL polystyrene FACS tube (Falcon, #352058) and centrifuged at 1200 rpm for 5 minutes at 4°C. We then lysed red blood cells for 15 minutes at room temperature using BD Pharm Lyse (BD, 555899). Cells were washed twice with staining media (Hanks balanced salt solution (HBSS) containing 2% FBS and 2 mM EDTA). The following antibodies were added (1:100) to each sample and incubated for 30 minutes at room temperature: Alexa Fluor 700 anti-mouse CD8a (Biolegend, 100730), PE/Dazzle-594 anti-mouse CD4 (Biolegend, 100456), APC anti-mouse CD19 (Biolegend, 115512), Alexa Fluor 488 anti-mouse NK-1.1 (Biolegend, 108718), PE anti-mouse CD3 (Biolegend, 100205) and Zombie Aqua Fixable Viability Kit (Biolegend, 423101) was used as a live-dead stain. We washed samples twice with staining media and sorted directly into TRIzol LS using a BD Aria FACS.

### qRT-PCR

We isolated and quantified RNA from sorted blood populations as described in the RNA Isolation and RNA-Seq Library Preparation. 100 ng of total RNA was used as input to generate cDNA using SuperScript IV VILO Master Mix (Invitrogen, 11756050), according to the manufacture protocol. cDNA was diluted 1:3 with DNase- and RNase-free water and 1 μL was used per each reaction. We performed qRT-PCR using FastStart Universal SYBR Green Master Mix with ROX (Sigma, 4913914001) on a ViiA 7 Real-Time PCR System (Thermo Fisher). Analysis was performed using the ΔΔCt method (Livak and Schmittgen, 2001). Primers used in qRT-PCR experiments are listed in the Sequences and Primers section.

### GEO Accession Numbers

All primary RNA-Seq data are available at the Gene Expression Omnibus (GSE124745 and GSE88819).

## SEQUENCES AND PRIMERS

### *In Situ* Hybridization riboprobe *-* mouse *Tug1* (492 bp)

GAGACACGACUCACCAAGCACUGCCACCAGCACUGUCACUGGGAACUUGAAGAUCCAAGUUUCUGUCCAGAACCUCAGUGCAAACUGACAACACUCCAUCCAAAGUGAACUACGUCCCGUGCCUCCUGAUUGCUGAAUGUUCACCUGGACCUGCCAAUGACCUUCCUUCUGCUACUCCAUCAGCCUACAGACCUGGUACUUGGAUUUUUGUCCAUGGUGAUUCCUUCCACCUUACUACUGAAGAAGACACCAUUCCAGUGGACCACUGUGACCCAAGAAGCAUUCAGCCAUCAUGAUGUGGCCUUUACCUCCACUCCUGUCCUACUCUGCCCAGAUUCAGCACAGCCCUUUAUAGUGCAGUCAAGAGUCUUCAAGCCAAAUAACUGAAGCUAUUUUAUCACAACAAAGGCCAGGUUUAUUCCAUAAAUGUACAGUUCAUUUCUGCAGUUUAUUCUUCAGAGACACAUAGUAAAUUUGGACCAGGGGAUUUUG

### Genotyping Primers

#### *Tug1*^*tm1.1Vlcg*^ knockout mice and MEFs

Tug1_5190-5166TD_Forward: TGACTGGCCCAGAAGTTGTAAG

Tug1_5190-5166TD_Reverse: GCAAGCAGGTCTGTGAGACTATTC

lacZ_5_Forward: TTGAAAATGGTCTGCTGCTG

lacZ_5_Reverse: TATTGGCTTCATCCACCACA

Ychr_Forward: TCTTAAACTCTGAAGAAGAGAC

Ychr_Reverse: GTCTTGCCTGTATGTGATGG

### Mouse *Tug1* (ENSMUST00000153313.2) cDNA clone

TTTTTTTTTTTTTTTTTTTTTTTTTTTTTGGGGGGGTTTTTTGTTTTGTTTTTTAAATTGAAGGCTAAAGTTTTTGAAAAAACTTTGTTGGACTCTGGCTGGGACACAAAATCAGATATTTGGAATCATTTTGAAGCTTAACTTTTTCCTAACCAGCCTTGTATTCTAATTGCTTGCAAATGTGAGACTGAATGGCCAAAATGCCGTTTGTTTGTTTGTTTATTGTCAGCTGCTTTTATCAAATTCCAGGCCATTATCCAGCAAACACTATTAAAATGTTTGAACAGTTGGGTTTCAAACATTTTTGTTTTGTGGAGTGGTGCTTATTAAGTGGTACAGCTCTCTAAGCAAGTGAACACAAACATATTTAAGTGTATTTTGTATGATTAGATGTTACCAATTCTGATATTTTATTCAAATGTCTAAAAAAATAAGTTGACTTATTCCCTTTACCAAAGGGCCAGAGACAAATGGTTTCCTTTCAAGAGAAATGACTGTTTTGAAGAAAAACTCTGTTGGTCTTAGCTCTTTTGTAATTAAATCTGGATGTACCTCAAAAGACTCTTTAAGACTGTGGTGTTAAAAGGCTTTCCTCTGGAGAAGGAGAAAAAATAAAATCAACTGGAACTTAAAAGCTTGAAATTCCATGACAAAACACAGATGTCCAGGATTGGAGGTTCATAAAGTACATGCAGTAGTTGGAGTGGATTCCATTTTCAGTGTAGCTGCCACCATGGACTCCAGGCTCCCAGATTTTCAAGAACTGGACCTGTGACCCAGAAGAGCTTGTCAAGATATGACAGGAACTCTGGAGGTGGACGTTTTGTATTCAATTTTGGAACTGTTGATCTTGCCGTGAGAAAAGAGAGACACGACTCACCAAGCACTGCCACCAGCACTGTCACTGGGAACTTGAAGATCCAAGTTTCTGTCCAGAACCTCAGTGCAAACTGACAACACTCCATCCAAAGTGAACTACGTCCCGTGCCTCCTGATTGCTGAATGTTCACCTGGACCTGCCAATGACCTTCCTTCTGCTACTCCATCAGCCTACAGACCTGGTACTTGGATTTTTGTCCATGGTGATTCCTTCCACCTTACTACTGAAGAAGACACCATTCCAGTGGACCACTGTGACCCAAGAAGCATTCAGCCATCATGATGTGGCCTTTACCTCCACTCCTGTCCTACTCTGCCCAGATTCAGCACAGCCCTTTATAGTGCAGTCAAGAGTCTTCAAGCCAAATAACTGAAGCTATTTTATCACAACAAAGGCCAGGTTTATTCCATAAATGTACAGTTCATTTCTGCAGTTTATTCTTCAGAGACACATAGTAAATTTGGACCAGGGGATTTTGTTTTGTTTATATTGTCAACACTGTCTGAAGAAAGGCATCTCTGAGAACAGCATTGGACCCTACTCCACAATCTCAAATGATTGAAGTTTCATAAACTGCCTAGGATCCTGTCAAGGCCACTGGACTCTTGTTCTTTTCCTACTTCAAAATCTGTAGCTGTCTACTAAATGACAAAGCAGATATTCTGACCCATTGGGATCAAAACCAAGGCATTTTGAATTCCTCATAGTATCATCTTCGGGTTACTCAGGAACCAAAACTTTTCACACCAATTTAAGAAATTCTACTGAGGAATCCCTTTACCTAACCATCTCACAAGGCTTCAACCAGATTCCTGAAAAGGCCTCTTGATATATCAAGATAGAACCTACATGCATTTTGTGAACAACTTATCACTGATTTTCCAAAGGCTTTGTGCTCTTGAAGTTCTTTGAAGGAAAGCTGTGTGGAAGTCCAGAGTAAAGTGAAGCTGCTCTGGATGAAGTAGTGAAGTGGGAGTTGAGGTCTACAACCTGCCACAACCATCTTCCTTTACCACCATGGTGATGCCAAAAGGGACTTCCTTAAAGCTCTTCAGAAAATCCTGCTTGAAACCACTACCCTAGACAACATGTTTGACCTGGATGGCATTCTCTTCAAAACAATTCATATTCAGTTGATGCTCAACATGTTTGGAGATGCTTTATTCAGAGAATGATGATAATTACAGCATTGTCTAATGAAGTTTTATTAATAGCATTCCATCCAAGGTGGACTTCCTGGAGTTGGATATAACCAGAGAGCAATTCATATGTATCCTACACTGAAGAACACCATTAACTTTCAGCAACCTATAGCTAGTGGTACTAGAAGTACGTGTCTTGGAAGTCTATGAGAGCTGGTATTGAAGCTGATGCCTCCTTAAGGCCATCTTAGACCAAGTTGTTTGTTTGACCTCTCCTCATTAACTATGGAGCAGAATTGAAATACAAATTTTTCCTAAAGGGACTTGCAACCTGGTTATCATTCATTATCTCAAGTTTCAAGTCATGTTGATGCAACCAGTAGTTATTAAACTGCTCCATGGTTTTTTGTTATTTAATACTTTTTCCAGGGCTTAAAAAAACAAAATTAAATTTCTCCAACACGTCTATACTTGTCTGTTCAAAAGTAACTACTCACCACTATATGGAACAGATGATTCTGAAGACACTCTGAGCATCCTTTATGATATTTGTGACTTAAAATGTGGCTGGAAATTTTCCTTCTACCCAGTGAAATATTTAATGATTAGTCTTCATGCCTGATACCATCAACTGTATATGCGTGATAGGCAAAGTTTGACATAGGCATTTGACTCTAGGCTATGATAGCTTGCTAGTAACTTCAAGTAGCATATTGTCAACCTGTTTGCTGGAAAAGTAGAGTAACTTGGAAAAAAAACTAAATGGCAGCTAAGGATTTTTTTCAGTATTCCTGAGTTTCTGTCCTTGGGATATTTCAATGAAATTTTCACCTGTCTCTTCACTTAACAGAGTGACTGACTCCTTACTATGAAGTATTCTTAAGACATTAAGATTACTTTTGTAGAAAGGATAAAATTCCTGACCATCCAAATCATCATAGTGAACAAGACTTCAATTTGTGACCTGAGAAAATCTCATTTCTCTACTTCGTAGTCAATGTAAGGGCCAATGCTATCAGCTACTCTGAGTGCACTGGGTAAACGTTGGAACTGCCTTCTTTATATCATTACTTTTTATCCTCTAAATTAATCATGGTTATGTAATTCTCGCCACAAATCAGCAAATCAGACTCAGATCTGGTTATTCTAGACTGCTCACAGTTAACAAATCAAACTCTGGATGACTTCTGCTTGTATATGCAACTACTATTTGTAAAGAAATTGCAAATTCACTTTTCTATTACCTCTACATTGCTAGCTCTTTCTTTTGTGTTTGTATTAAAAACAAAAATAAGCTACACTGCCAGCTATTCCCTCCTGCCATACTCAGTTAAAATGAAGAATCGGGAATCTAACCAGTGAATGGATAAGTAGAAAAAACTAAAACTTAAGGCAAAAGCCTTAATCTAGGGCCTTTTCTACTATCTTCATGTCTTGGATTTCATCTAAAATCAACAGTGCCACCCAACCAGTCTGAGGTCTTGACTTGCTTTTAAGATGATTCTTAGAGATGGGCTGTATTACAGAAGGTGAAGACTTGATTACCAAAGAAAGTAGAGCCAACTTTGACAAACCTGGCTCTACAATCCTATTGCTTCCAGATGTAGCATAGACTCATAACTAGAACCTCAAGTCTGCATTGAGGATATAGCCTTCTAAGCTGACAGTTCTTGCAACAGGTGAGCAAGAAAATGAAAGCTGTTATACCCAACTGGCCCTTTAAGATCCAAAAATAATGTCTGGACTAAACCCTATGGAGTACCCAGGACAAAAACTAATTTACAGAGCTTCATTATTAATCTGCCTGTTCTTCTAGCTTAATTATTGGTATGGCTGGCCCTACTGAAGTAGTTTGTCTGTTTACCTGTCTTCAGCTCTTAACCTGGCTATTTTGACATGCTACTGCAATTAGACTAACTGGCTTTGAGAAGACTACAATCAGTTTCAGCCTCTCCTTTGCCCAATTTCACCAAGGAATTTTGATAAGAGGAACCCATACCTCACCCCACCAGAACAGAAAGGACCATGCTGCATATTCCTTGACCAGCAACTTTAAGTAGAGAACAACCCTGCTTGTTTTCAACATCTGAAACACCATTTGATCTAATAGGAGTATAGAAGGTTGACAGCAGAGTACACTACTTACTTCTTTCATAACTCAGAAATGAATATGACTGGCCCAGAAGTTGTAAGTTCACCTTGACAAGAAACAGCAACACCAGAAGTTTACTGCTGAACTTAACTTGCCACTTACTCGAATAGTCTCACAGACCTGCTTGCCAAGTAGGAGGCTAGTTTTCCTGCTTCATATCACCATTGGAGTGGGGCTCAATGGGGTCAATGTTAATACTGACTTGAATGGGGACCTTATGGTGAATCCTAGACTATGAGGCTAATGGAAATTATTGTCTATTCAAGTGGATTATAGATTTCCTGAGGACAGAACAGACATCACTCCTGGTGATTTTTAGAACTTGATTACCAAGGAAGAAATACCAGCTGCTAACAGTCAACTTCATGGGCAAAGATTAAGCTCTCTATATCTGGTCGTATCCTGGATGCTAGTTTTTTATTGCCCAGTGACCATTTCCATCTCACGCTTAACTTCCTGATGTTTTTTGGAACCATCTCTTCCAATTTTCAGTCCTGGTGATTTAGACAGTCTTTTCATGCTGGACATTTTGTTGCAACCTCATCAATCACAGCAAAGTCCATCTTGACTTTAGTGATTAGTTCAGGAATGGATGCATGATTCAAGTTTGTCCAATGATAATCAACCCTAGGTGTTTTCTCAGTTGTGGAGAAGTTCTCTTAGATGCTTTAGCTTTGTAGGAGAAAACTCAAACCAACAGGGCCTACCTACTATGTTGAATGATTGTAGGAGAAAACTCAAACCAACCAGGCCTACCTACTATGTTGAATGAGCCAGGCAGAAAATGAAGCCAGTACAGAGGGAAATGGAGCCAAAAGAGGAAGAGACTTGAGTTCTGATGATCACATTTATGCCCCTGTATCCAACTGTGCCTGAAGCTAATAGTACATCACCTGGACTTTTCAGTTATGTGAACCAATAAATTCCCCTTTTTGTTTAAGTTACTTTGAGTT

### Full Length *Tug1* primers for cloning in pTRE2pur

#### Tug1_MluI_cDNA_Fwd

gagaacgcgtTTTTTTTTTTTTTTTTTTTTTTTTTTTTGGGGGGGTTTTTTGTTTTGTTTTTTAAATTGAAGGCTAAAGTTTTTGAAAAAACTTTGTTGGACTCTGGCTGG

#### Tug1_EcoRV_cDNA_Rev

gagagatatcAACTCAAAGTAACTTAAACAAAAAGGG

### Primers for full length sequencing of pTRE2-*Tug1* expression vector

LNCX AGCTCGTTTAGTGAACCGTCAGATC

TUG1_76 TTTAAATTGAAGGCTAAAGTTTTTGAA

TUG1_266 GGCCATTATCCAGCAAACAC

TUG1_755 ACTCCAGGCTCCCAGATTTT

TUG1_1254 TCTTCAAGCCAAATAACTGAAGC

TUG1_1741 AGAACCTACATGCATTTTGTGAA

TUG1_2268 ATGCCTCCTTAAGGCCATCT

TUG1_2754 TGTCAACCTGTTTGCTGGAA

TUG1_3267 TTGCAAATTCACTTTTCTATTACCTC

TUG1_3746 CCCAACTGGCCCTTTAAGAT

TUG1_4241 TGACAAGAAACAGCAACACCA

TUG1_4740 TCACAGCAAAGTCCATCTTGA

### Primers for qRT-PCR

Tug1_Fwd CTCTGGAGGTGGACGTTTTGT

Tug1_Rev GTGAGTCGTGTCTCTCTTTTCTC

Gapdh_Fwd GGTGAAGGTCGGTGTGAACG

Gapdh_Rev CTCGCTCCTGGAAGATGGTG

### Primers for sub-cloning full length *Tug1* from pTRE2-*Tug1* into pcDNA3.1(+) expression vector

Tug1_Tg F/KpnI ataggtaccGCCCCGAATTCACGCGTT

Tug1_Tg R/NotI atagcggccgcACCTGAGGAGTGAAGA

### Human TUG1-BOAT (ORF1) sequences with 3xFLAG (blue) for expression construct design

#### Human ORF1

CTGGCGCGCCCTCCCCCCCTCCCGGGTCTGGTAGGGCGAAGGAACGGGCGTGCGGTCGATCGAGCGATCGGTTGGCGGCTCTTTCTCCTGCTCTGGCATCCAGCTCTTGGGGCGCAGGCCCGGCCGCCGCGGCGCGCGCCCGGTGGCCGTTGGCGCTCGCGCCGCGTCTTTCTTCTCGTACGCAGAACTCGGGCGGCGGCCTATGCGTTTGCGATTCGACGAGGAGTCGTCCGGGTGGTCGGCGGCGGCGGGCAGCTGCTCCGCCCCGCTCCGGGGGAGGCGGCGGCGGCAGCGGCCGCGGGATTTGGAGCGGCCGGGGAGGCGGGGGTGGCCGGGGCCGGCTTGGAGGCCTGGCGCCACCCTTCGGGGCCTGCAAGGACCCAGTTGGGGGGGCAGGAGGGGGCCGGAGGATGGTTGGTTGTGGGATTTCTACTTTGCCTTTTCCTCCTTATGCCGCCTGACTACAAAGACCATGACGGTGATTATAAAGATCATGACATCGACTACAAGGATGACGATGACAAGTAG

#### Human ORF1+UTR

GGCCGAGCGACGCAGCCGGGACGGTAGCTGCGGTGCGGACCGGAGGAGCCATCTTGTCTCGTCGCCGGGGAGTCAGGCCCCTAAATCGAAGAAGCCCTGGCGCGCCCTCCCCCCCTCCCGGGTCTGGTAGGGCGAAGGAACGGGCGTGCGGTCGATCGAGCGATCGGTTGGCGGCTCTTTCTCCTGCTCTGGCATCCAGCTCTTGGGGCGCAGGCCCGGCCGCCGCGGCGCGCGCCCGGTGGCCGTTGGCGCTCGCGCCGCGTCTTTCTTCTCGTACGCAGAACTCGGGCGGCGGCCTATGCGTTTGCGATTCGACGAGGAGTCGTCCGGGTGGTCGGCGGCGGCGGGCAGCTGCTCCGCCCCGCTCCGGGGGAGGCGGCGGCGGCAGCGGCCGCGGGATTTGGAGCGGCCGGGGAGGCGGGGGTGGCCGGGGCCGGCTTGGAGGCCTGGCGCCACCCTTCGGGGCCTGCAAGGACCCAGTTGGGGGGGCAGGAGGGGGCCGGAGGATGGTTGGTTGTGGGATTTCTACTTTGCCTTTTCCTCCTTATGCCGCCTGACTACAAAGACCATGACGGTGATTATAAAGATCATGACATCGACTACAAGGATGACGATGACAAGTAG

### Mouse TUG1-BOAT (ORF1) sequences with 3xFLAG (blue) and HA (red) tags for expression construct design

#### Mouse ORF1

CTGGCGCGCCCTCCCCCCCTCCCCGGTCTGGTAGGGCGAAGGAGCGGGCGTGCGGTCGATCGAGCGATCGGTTGGCGGCTCTTTCTCCTGCTCTGGCATCCAGCTCTTGGGGCGCAGGCCCGGCCGCCGCGGCGCGCGCCCGGTGGCCGTTGGCGCTCGCGCCGCGTCTTTCTTCTCGTACGCAGAACTCGGGCGGCGGCCTATGCTTTTGCGATCCGACGAGGGGTCGTCCGGGTGGTTGGCGGCGGCGGGCAACTCCGCCCCGCTCCCGGGGAGGCGGCGGGGGAAGCTGGGGTGGCCGGGGCTGGCCTGGAGGCCTGGCGCCACCCCTCGGGGCCTGCTAGGACCCAGTTGGAGGGTCAAGAGGGAGCTGGAGGATGGTTGGTGGTGGGCTTCCTCCTTTGCCTTTTCCTACTTATGCCACCTGACTACAAAGACCATGACGGTGATTATAAAGATCATGACATCGACTACAAGGATGACGATGACAAGTACCCATACGATGTTCCAGATTACGCTTAG

#### Mouse ORF1+UTR

GGCCGAGAGACGCAGCCGGGACGGTAGCTGCAGAGCAGAGCGGAGGAGCCATCTTGTCTTGTCGCCGGGGAGTCAGGCCCCTAACTCGAAGAAGCCCTGGCGCGCCCTCCCCCCCTCCCCGGTCTGGTAGGGCGAAGGAGCGGGCGTGCGGTCGATCGAGCGATCGGTTGGCGGCTCTTTCTCCTGCTCTGGCATCCAGCTCTTGGGGCGCAGGCCCGGCCGCCGCGGCGCGCGCCCGGTGGCCGTTGGCGCTCGCGCCGCGTCTTTCTTCTCGTACGCAGAACTCGGGCGGCGGCCTATGCTTTTGCGATCCGACGAGGGGTCGTCCGGGTGGTTGGCGGCGGCGGGCAACTCCGCCCCGCTCCCGGGGAGGCGGCGGGGGAAGCTGGGGTGGCCGGGGCTGGCCTGGAGGCCTGGCGCCACCCCTCGGGGCCTGCTAGGACCCAGTTGGAGGGTCAAGAGGGAGCTGGAGGATGGTTGGTGGTGGGCTTCCTCCTTTGCCTTTTCCTACTTATGCCACCTGACTACAAAGACCATGACGGTGATTATAAAGATCATGACATCGACTACAAGGATGACGATGACAAGTACCCATACGATGTTCCAGA TTACGCTTAG

## DECLARATIONS

### Ethics Approval and Consent to Participate

Not applicable

### Consent for Publication

Not applicable

### Availability of Data and Materials

The datasets generated and/or analyzed during the current study are available in the Gene Expression Omnibus repository, GSE124745 and GSE88819.

### Authors’ Contributions

Conceptualization, M.S., J.L.R.; Methodology, M.S., E.J.P., J.L., G.D., T.H.; Computational Analyses, T.H., A.F.G., N.D.R.; Experimental Analyses, M.S., E.J.P., A.F.G., N.D.R., J.L., G.D., A.W., C.G., C.K., K.T.; J.S., C.M., S.H.; Investigation, M.S., E.J.P., S.C.L., C.G., J.L., G.D., A.W., N.C., C.K., K.T., J.S., C.M., S.H.; Data Curation, M.S., E.J.P. T.H., J.L., G.D., A.W.; Writing Draft, M.S., J.L.R., G.D., J.L., C.M. T.H.; Figures, M.S., E.J.P., C.M., J.L., G.D., A.W., T.H.; Supervision, Administration, and Funding Acquisition, J.L.R., H.E.H., N.H.; All authors have read and approved the final manuscript.

#### Acknowledgements

We thank Diana Sanchez-Gomez for assistance in the mouse facility; Catherine MacGillivray and Diane Faria in the HSCRB histology core facility; Joyce LaVecchio and Nema Kheradmand in the HSCRB flow cytometry core; and the Broad Institute of MIT and Harvard Genomics Platform and the Bauer Core Facility at Harvard University for RNA-sequencing support. We also thank Dr. James C. Lee and Philipp Maas (PhD) for manuscript feedback.

## Authors’ Information

Not applicable

## SUPPLEMENTARY FIGURES

**Figure S1.**
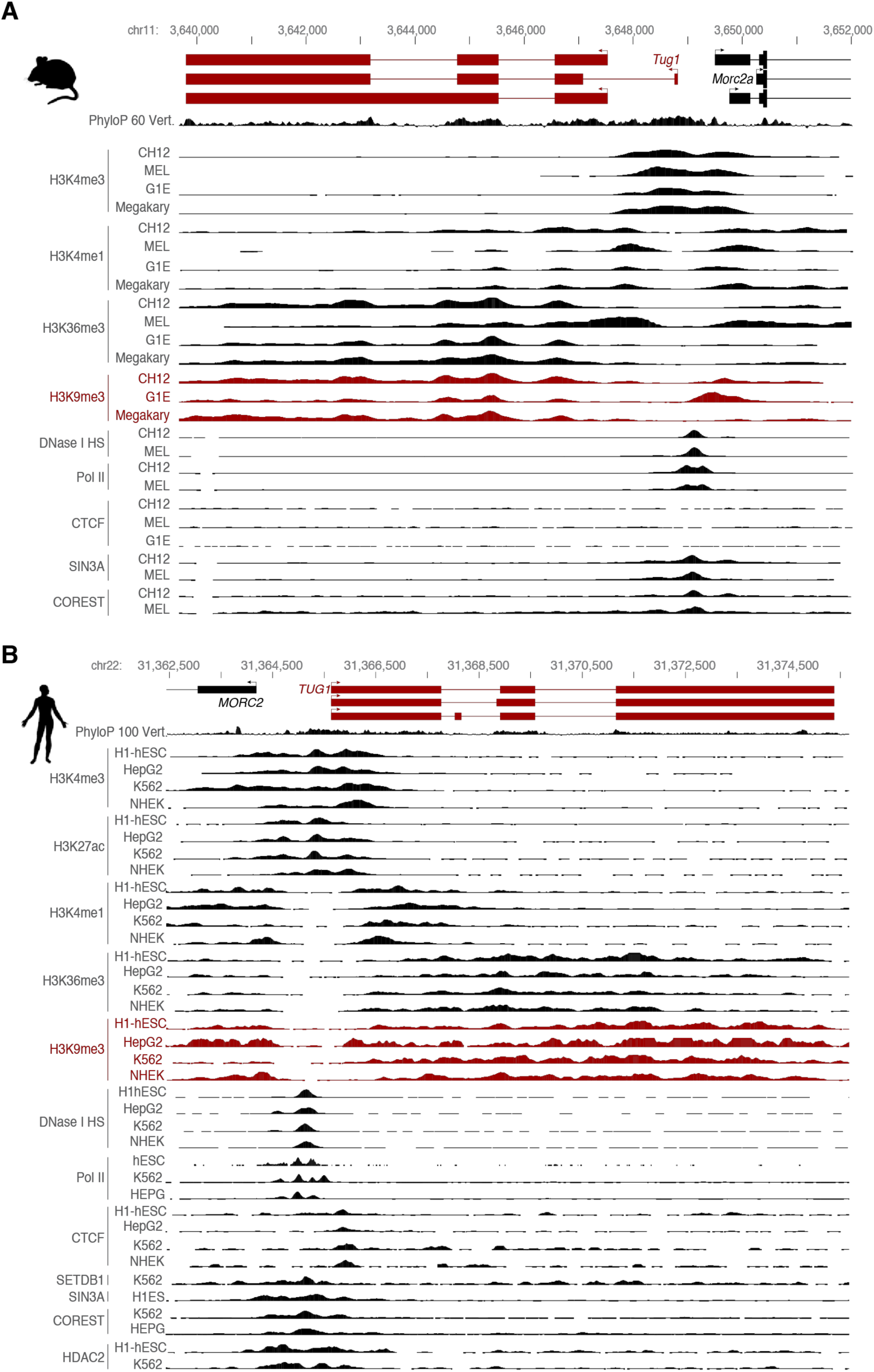
Mouse and human *Tug1* locus and chromatin context in different cell types. **(A)** *Tug1* mouse and **(B)** human genomic loci. Evolutionary nucleotide conservation (PhyloP) of the locus are presented along with the chromatin context (DNase I hypersensitive regions, histone modifications) and protein binding ChIP-seq peaks (Pol2, CTCF, SIN3A, COREST, SETDB1, HDAC2) from ENCODE (UCSC Genome Browser, mm9) datasets in the indicated cell types.

**Figure S2.**
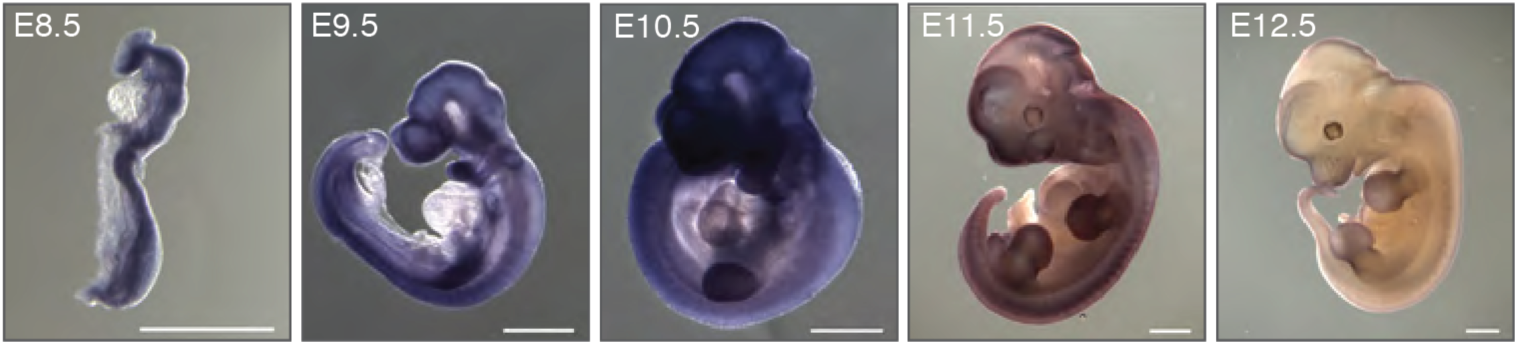
*In vivo* expression pattern of *Tug1* during murine embryogenesis. RNA *in situ* hybridization of *Tug1* RNA using a digoxigenin-labeled antisense RNA probe in mouse embryos at different developmental stages. Embryonic day (E)8.5, E9.5, E10.5, E11.5, and E12.5 are shown.

**Figure S3.**
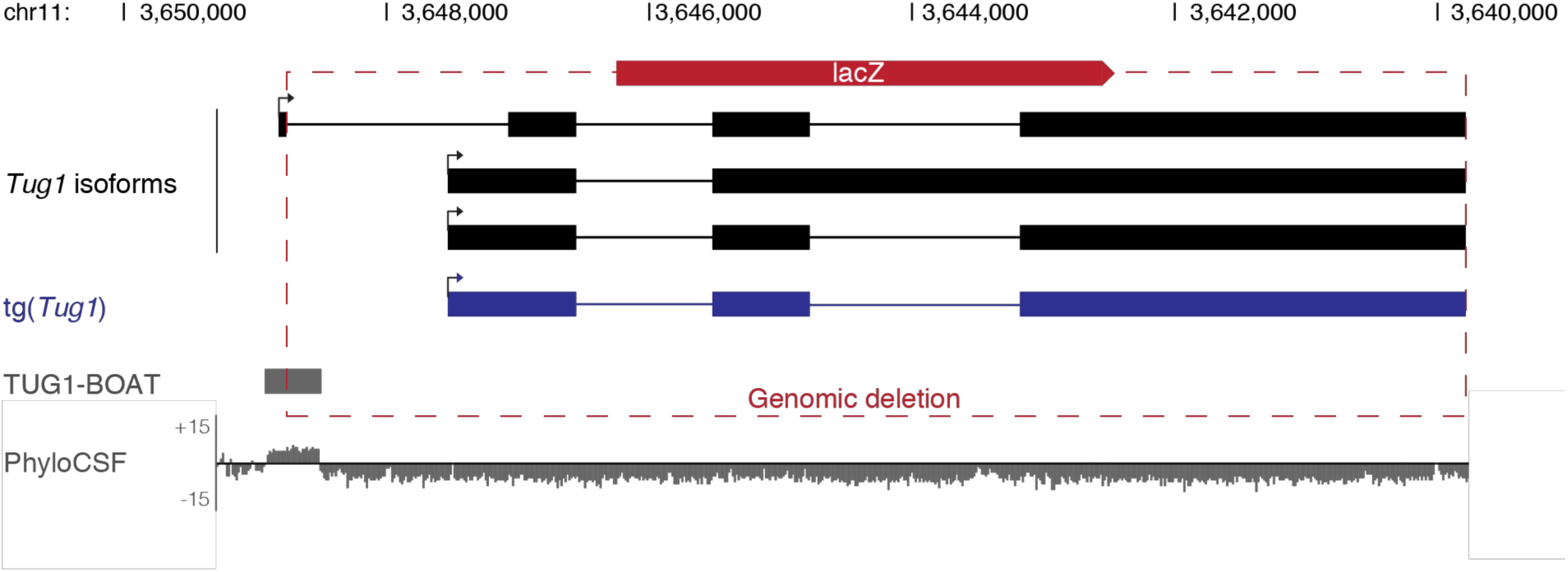
Overview of the *Tug1* locus in mouse. UCSC genome browser showing the murine*Tug1* locus. The three predominate *Tug1* isoforms are depicted (black) and the *Tug1* transgene (tg(*Tug1*)) is shown (blue). For *Tug1* knockout, the longest annotated *Tug1* isoform was replaced by a *lacZ* reporter cassette, leaving the promotor and first exon intact. The deleted region is indicated by red dashed lines. The open reading frame (ORF) encoding the TUG1-BOAT peptide and PhyloCSF scores for the (−2) frame across the locus are depicted (grey). Chromosomal coordinates (mm10) are shown.

**Figure S4.**
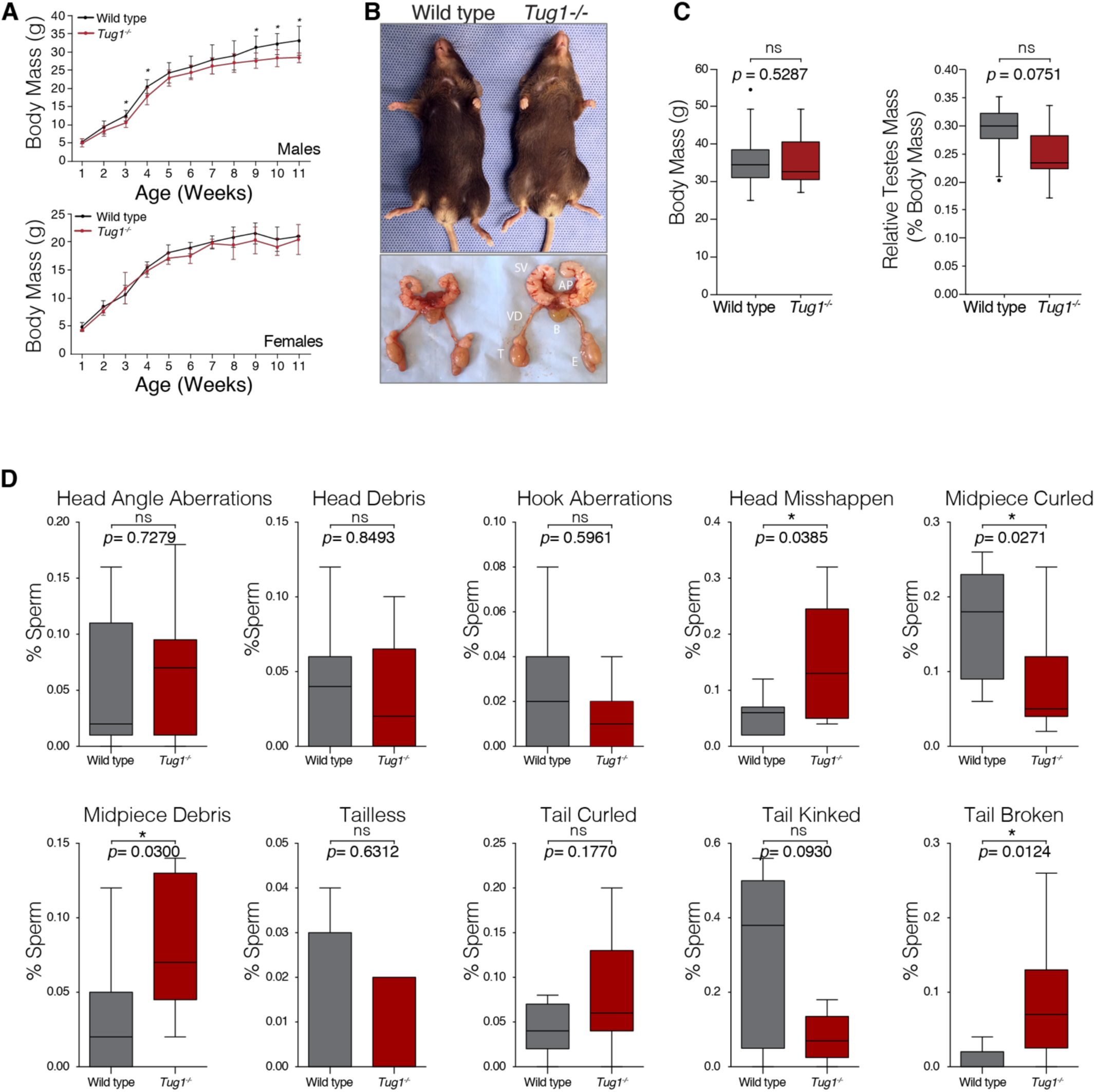
Morphology analysis of *Tug1*^-/-^ mice and sperm. **(A)** Body mass (g) measurements over 11 weeks of male and female *Tug1*^-/-^ mice compared to wild type littermates. Males: *Tug1*^-/-^ (n = 7); WT (n = 8). Females: *Tug1*^-/-^ (n = 3), WT (n = 7). Significant p values at specific time points are indicated (*). **(B)** Representative images from adult male mice (12 weeks old) show normal physiological appearance of external genitalia and reproductive tracks in *Tug1*^-/-^ compared to WT. Seminal vesicles (SV), vas deferens (VD), bladder (B), testicle (T), epididymis (E), anterior prostate (AP). **(C)** Box plots of body mass (g) (left panel), relative testis mass (testis mass / body mass; middle panel) and total sperm count for wild type (n = 9) and *Tug1*^*-/-*^ males. **(D)** Box plots of the percentage of different sperm morphological abnormalities for wild type (n = 9) and *Tug1*^-/-^ (n = 8) males. Significant (*) p value (Wilcoxon rank sum test) is indicated.

**Figure S5.**
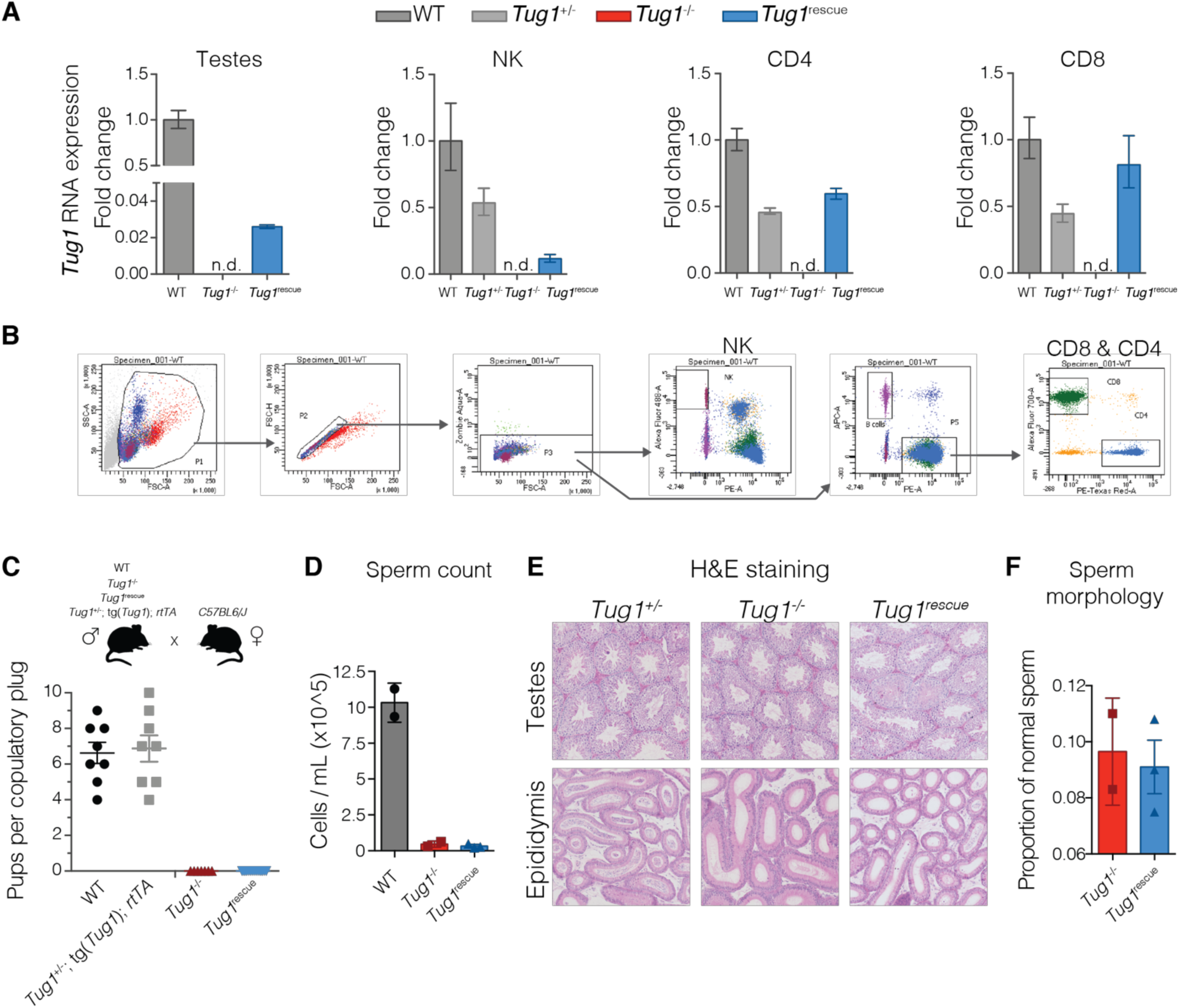
*Tug1* transgene expression and fertility assessment. **(A)** qRT-PCR for *Tug1* RNA expression in testes and sorted peripheral blood populations: WT (*n* = 1), *Tug1*^+/-^ (*n* = 1), *Tug1*^-/-^ (*n* = 1), and *Tug1*^rescue^ (*n* = 1) and sorted peripheral blood populations. Error bars indicate the relative quantification minimum and maximum confidence interval at 98%. Not detected (n.d.). **(B)** Representative flow cytometry gating strategy for NK, CD4, and CD8 cells in peripheral blood from WT, *Tug1*^+/-^, *Tug1*^-/-^, and *Tug1*^rescue^ mice (gating from WT peripheral blood shown). **(C)** Scatter dot plot (mean with standard error of the mean shown) of the number of pups at birth per copulatory plug for matings using male wild type, *Tug1*^*+/-*^; tg(*Tug1*); rtTA, *Tug1*^*-/-*^, or *Tug1*^rescue^ (on dox diet) with wild type C57BL/6J females. Each dot represents a litter from a different mouse. **(D)** Sperm count from control (WT and *Tug1*^+/-^, n = 2), *Tug1*^-/-^(n = 2), and *Tug1*^resuce^ (n = 3) mice. Each dot represents a different mouse and the error bars indicate the standard error of the mean. **(E)** Hematoxylin and eosin staining in *Tug1*^+/-^, *Tug1*^-/-^, and *Tug1*^rescue^ testes and epididymis. **(F)** Morphological analysis of sperm from *Tug1*^-/-^ (n = 2), and *Tug1*^rescue^ (n = 3) mice.

**Figure S6.**
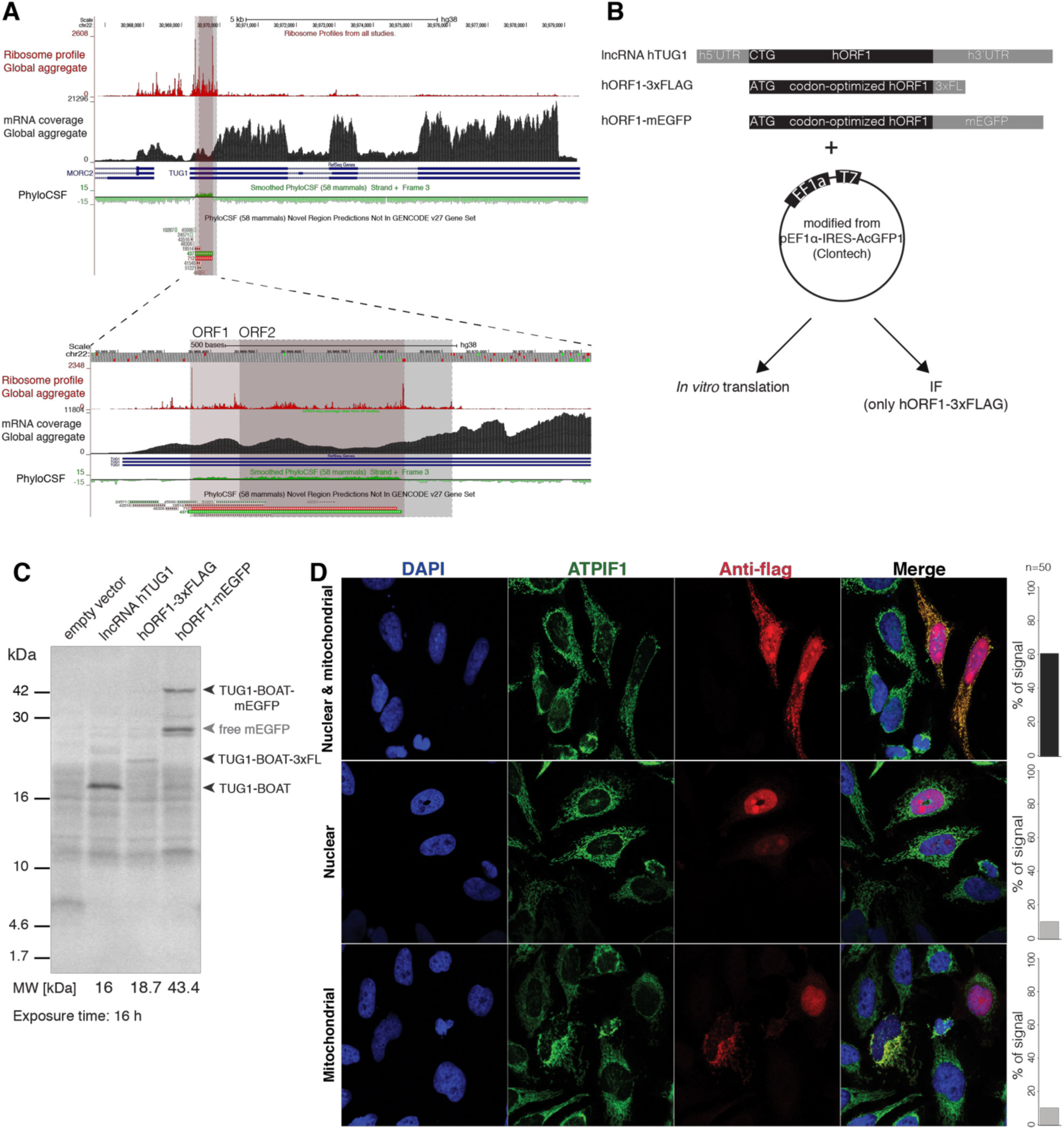
The 5’ region of human *TUG1* encodes a conserved peptide. **(A)** GWIPS-viz tracks for human *TUG1* genomic locus (hg38) is shown. Global aggregate of ribosome occupancy (ribosome profile), RNA-seq (mRNA coverage), and evolutionary protein-coding potential (PhyloCSF) across the *TUG1* locus is shown. ORF1 and ORF2 are outlined with red and gray boxes, respectively. Tracks surrounding both ORFs are zoomed in for clarity (bottom). **(B)** Scheme of additional human ORF1 construct design. hORF1 was left unlabeled with its endogenous non-canonical start codon (CUG) and placed between its native 5’ UTR and 321 nucleotides of its 3’ UTR (lncRNA h*TUG1*). hORF1 was codon optimized to contain the canonical AUG start codon and labeled with either a 3xFLAG epitope tag (hORF1-3xFLAG) or mEGFP (hORF1-mEGFP), inserted prior to the stop codon. Constructs were inserted into a modified form of pEF1α-IRES-AcGFP1 and assessed for *in vitro* translation (shown in C). hORF1-3xFLAG was additionally analyzed by immunofluorescence (IF) (shown in D). **(C)** The synthesis of peptides from all three constructs and an empty vector control was assessed using a wheat germ extract *in vitro* translation assay. Newly synthesized peptides are labeled with arrows and correspond to their respective predicted molecular weights (16 kDa for hTUG1-BOAT, 18.7 kDa for TUG1-BOAT-3xFLAG, and 43.3 kDa for TUG1-BOAT-mEGFP). **(D)** Localization of codon-optimized 3xFLAG tagged ORF1 (hORF1-3xFLAG) was assessed by immunostaining against the 3xFLAG (red) in HeLa cells. Nuclear localization was monitored by DAPI (blue) and mitochondrial localization was monitored by the organelle marker ATPIF1 (green).

## SUPPLEMENTARY TABLES

**Table S1.** *Tug1*^-/-^ and wild type sperm morphological defects.

**Table S2.**Prostate, spleen, eyes, brain, heart, liver, and MEF RNA-seq.

**Table S3.** Allele-specific RNA-seq in testes.

**Table S4.** Testes RNA-seq and *Tug1*^rescue^ RNA-seq in testes.

All supplementary tables are available at the Gene Expression Omnibus (GSEA124745 and GSE88819).

